# Frontal Brain Injury Reduces Sensitivity to Reward-Predictive Cues and Remodels the Nucleus Accumbens

**DOI:** 10.64898/2026.04.17.718474

**Authors:** Erskine Chu, Jenna E. McCloskey, Mia A. Eleid, Sathvik Jami, Alexandra G. Dorinsky, Fikir B. Arega, Kris M. Martens, Fangli Zhao, Jonathan M. Packer, Patrick Stevens, Maciej Pietrzak, Candice C. Askwith, Jonathan P. Godbout, Cole Vonder Haar

**Author notes:** Corresponding author: Cole Vonder Haar, Department of Neuroscience, Ohio State University 460 W 12^th^ Ave, Columbus, OH 43210. These authors contributed equally (Erskine Chu, Jenna E. McCloskey).

## Abstract

Traumatic brain injuries (TBIs) are more than mere lesions and generate a persistent secondary pathology. This, combined with functional reorganization of circuits post-injury, may explain the increased risk for psychiatric disorders in patients with TBI. In the current studies, we demonstrate that frontal TBI changed the Pavlovian behavioral response to reinforcer-predicting cues and reduced the motivational value of cues. TBI also chronically impaired decision-making on a gambling-like task with reinforcer-paired cues. To investigate how these changes occur, we evaluated the nucleus accumbens (NAc) core. At a subacute time point (14 days), we confirmed reduced input to the NAc with optogenetics and evaluated electrophysiological and transcriptional changes. TBI increased neuronal excitability and the single nucleus RNA sequencing profile indicated a substantial stress and inflammatory response, but also high indicators of plasticity, particularly in D1- and D2-positive medium spiny neurons. To evaluate how these subacute changes transitioned to chronic NAc dysfunction, we measured immunohistochemical surrogates of activity post-mortem and recorded calcium activity from the NAc after TBI during Pavlovian conditioning. TBI reduced histological markers of activity and reduced cue-evoked calcium activity. Overall, these data indicate that substantial reorganization of the NAc occurs following frontal brain injury. A primary effect of this is to reduce the salience of environmental cues linked to outcomes. The inability to properly process outcomes could contribute to broader psychiatric symptoms after TBI, including impairments in decision-making, behavioral flexibility, and impulsivity but also presents a potential treatment target.

## Introduction

Among the types of insults which can alter brain function, traumatic brain injuries (TBIs) are one of the most drastic. The initial impact can immediately disrupt brain circuits (e.g., loss of consciousness) via direct cell death and strain to axons. More insidious is the wide-scale secondary pathophysiology cascade which can have effects on the scale of minutes and hours (e.g., excitotoxic cell death) all the way up to years (e.g., neuroinflammation and neurodegeneration) [1, 2]. Even relatively local injuries influence distal circuits due to loss or dysfunction in projections, resulting in a progressive pathology at distal locations [3]. The resulting adaptation and reorganization in response to this process may contribute to the increased incidence of psychiatric disease after TBI.

Clinically, brain injury increases risk for a variety of psychiatric symptoms, including impulsivity, risky decision-making, and poor behavioral flexibility [4–8]. Given that these symptoms can involve impaired learning from salient environmental consequences (i.e., reinforcement and punishment), this may explain the increased comorbidities of conditions such as gambling disorder [9, 10] and substance use disorders [11] in patients with TBI. Non-remitting clinical symptoms are also replicated in rats after frontal TBI [12–15], providing a means to understand mechanisms of psychiatric disorder risk. However, the interplay between loss of tissue, functional reorganization, and ongoing pathology is not well understood in the context of these disorders. Brain injury may also alter fundamental reinforcement-related behaviors. Our group previously identified a surprising outcome when examining simple Pavlovian conditioning after TBI in rats. Pavlovian conditioning generates sign-tracking (orienting/attending to stimuli predicting reinforcement) and goal-tracking (orienting toward the location of the reinforcer) behaviors. For intact rats, these behaviors are distributed fairly uniformly [16], however after TBI, rats showed near-exclusive goal-tracking behavior [17]. This fundamental change to engagement with reinforcer-predictive cues in the environment may reflect a broader inability to efficiently integrate environmental feedback (e.g., a choice resulting in suboptimal outcomes), adjust behavior, and maximize reinforcement.

The nucleus accumbens (NAc) is a critical structure involved in both Pavlovian cue processing and broader decision-making. The NAc is highly integrative, receiving convergent input from multiple regions, including the ventral tegmental area (VTA), medial prefrontal cortex (mPFC), basolateral amygdala, ventral hippocampus, and thalamus [18]. For Pavlovian learning, dopaminergic projections of the VTA to NAc have been a primary circuit of study. Inhibition of these projections at the time of cue presentation attenuated development of a sign-tracking response [19]. In operant choice tasks, the NAc is essential to invigorate responses likely to result in reinforcement and is critical for learning and development of preference. Pharmacologic inactivation of the NAc core and a broad dopamine receptor antagonist suppressed the ability to distinguish beneficial probabilistic outcomes [20, 21]. Together, these indicate the NAc as a core hub for integrating probabilistic information and Pavlovian learning.

In the case of TBI, the NAc may serve as a key node for many of the dysfunctional behaviors described above. Clinical studies demonstrated marked volumetric or structural change in the NAc in populations with a history of TBI [22–25]. Moreover, deep brain stimulation of the NAc for severe TBI successfully reduced symptoms (including impulsive decisions) in a small number of cases (N = 5) [26, 27]. Despite clinical relevance, animal studies have not prioritized linking psychiatric symptoms and NAc function after TBI. Reconciling fundamental findings on NAc function with clinical conditions like brain injury is of the utmost importance so that animal models can be used to efficiently screen for future therapeutics.

Thus, the goal of the current study was to test the hypothesis that frontal TBI alters information processing in the NAc, impairing processing of reinforcer-associated cues and impairing behavior. This was accomplished by validating TBI-induced behavioral phenotypes and their link to the NAc on two distinct tasks involving reinforcer cues (reinforcer-predictive vs. reinforcer-paired). We then probed projection loss using optogenetics and examined the consequences of this loss on cellular physiology and transcriptional profiles using single nucleus RNA sequencing. Finally, we validated impaired NAc processing of reinforcer-predicting cues in real time using fiber photometry calcium imaging. To our knowledge, this is the first report of cell-level sequencing of the accumbens and on-task calcium imaging in a model of TBI, providing a strong dataset for linking TBI to the many other disease models using these methods.

## Results

Results for several variables not presented here can be found in the supplement. Supplement 1 contains additional figures and analysis. This includes a breakdown of behavioral measures by sex (Fig S1, S2, S3, S5). Supplement 2 contains spreadsheets with full statistical tables for all analyses and full pathway lists for sequencing data.

### TBI reduced orienting to and motivation for reinforcement-predictive cues

Rats received a TBI to the mPFC and then were tested at 10 days post-injury in a visual Pavlovian conditioning task [17] which randomly presented one of two levers and the light above that lever (Fig 1A). One lever-light pair predicted a sucrose pellet reinforcer after 5 s (CS^+^) and the other was never associated with reinforcement (CS^-^). “Sign-tracking” behavior was recorded as duration of lever pressing as a proxy for interaction; “goal-tracking” behavior was recorded as duration of time in the food hopper. TBI significantly reduced the ratio of sign-tracking to goal-tracking behaviors for the CS^+^ (*p* < 0.001; Fig 1B). When re-tested at a chronic time point (98 days post-injury), these behavioral differences persisted (*p* < 0.001; Fig 1B). TBI also increased the time it took to discriminate CS^+^ from CS^-^ cues (*p* < 0.001; Fig S2A).

**Figure 1.**
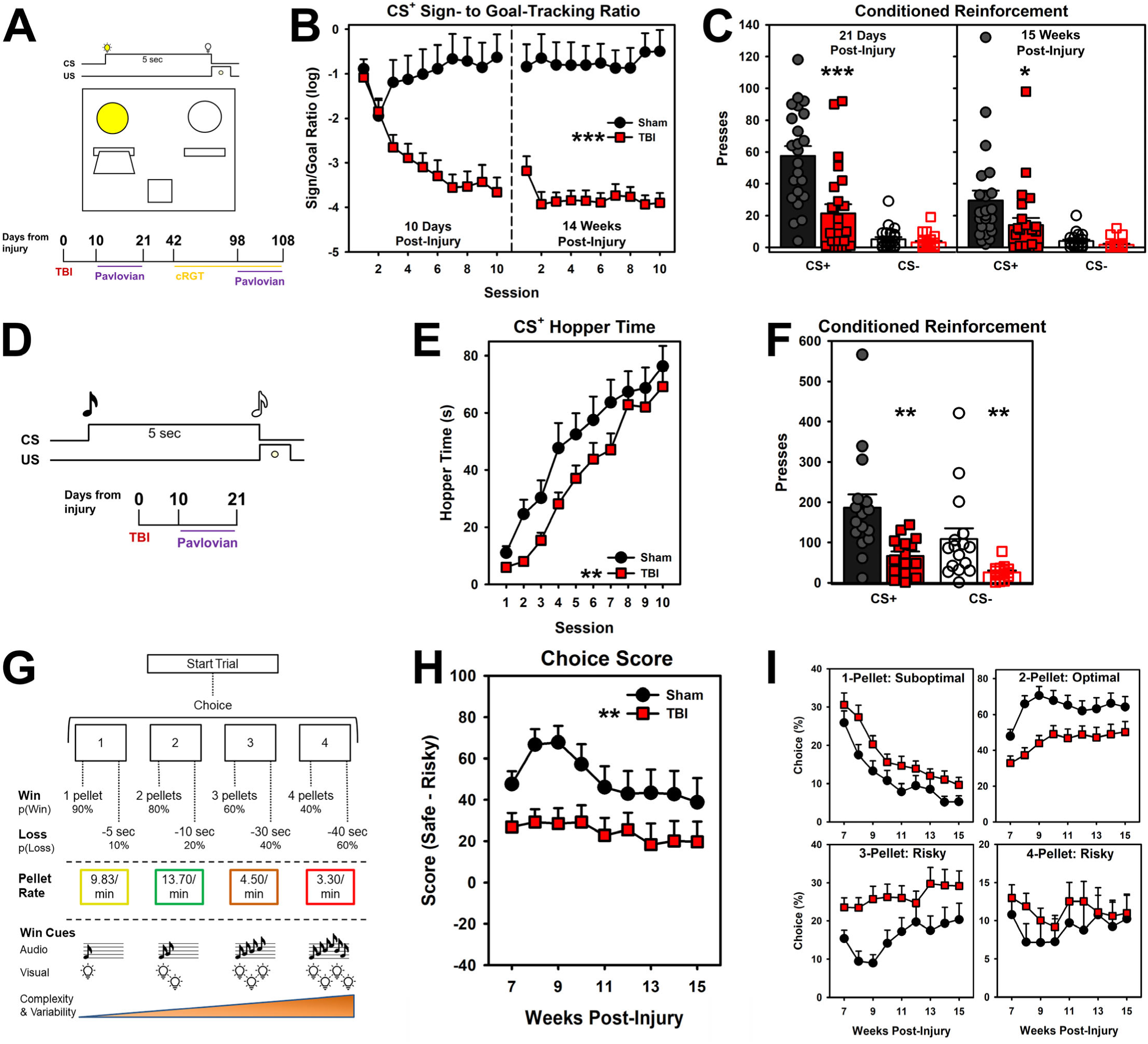
TBI reduced orienting to and motivation for reinforcement-predictive cues and decreased optimal decision-making on a cued rat gambling task (cRGT). Rats were subjected to TBI or sham procedures and then assessed on visual Pavlovian conditioning, cRGT, then again on visual Pavlovian conditioning (N = 47; panels A-C, G-I) or on auditory Pavlovian conditioning (N = 32; panels D-F). A) Schematic of the visual Pavlovian conditioning task showing a CS^+^ trial in which a lever & light cue were presented preceding a sucrose pellet (CS^-^ trials used opposite lever and no pellet) and timeline of experiments for data in panels B, C, H, and I. B) Sign-to-goal-tracking ratio for the CS^+^ at acute (10 days post-injury) and chronic (14 weeks post-injury) time points. TBI shifted behavior toward goal tracking at both time points (*p*’s < 0.001). C) Number of presses to receive the cue alone on a conditioned reinforcement probe immediately following Pavlovian conditioning in the same rats as panel B. TBI reduced responding for the CS^+^ cue as a conditioned reinforcer at both time points (acute: *p* < 0.001, chronic: *p* = 0.05). D) Schematic of the auditory Pavlovian conditioning task showing a CS+ trial in which an auditory cue was presented preceding a sucrose pellet (CS^-^trials used a different tone and no pellet) and timeline of experiment for data in panels E-F. E) Responding in the hopper during CS^+^ presentation. TBI initially reduced hopper responding but approached sham levels by the end (*p* = 0.001). F) Number of presses to receive the cue alone on a conditioned reinforcement probe immediately following Pavlovian conditioning in the same rats as panel E. TBI reduced responding for the CS^+^ cue as a conditioned reinforcer and the CS^-^ cue (*p* = 0.002). G) Schematic of the cued Rodent Gambling Task (cRGT), showing four choice options with distinct reinforcement, punishment, and cue complexity profiles. H) Overall risk preference score on the cRGT. TBI reduced score, indicating a shift toward risk preference. I) A breakdown of individual choice options on the cRGT that make up the choice score. TBI reduced preference for the optimal (2-pellet) option in favor of suboptimal (1-pellet) and risky (3-pellet) options. Data are means + SEM with individual subject data superimposed as points on top of bars; *** = *p* < 0.001, ** = *p* < 0.01, * = *p* < 0.005.

To evaluate if reinforcer-predicting cues were inherently motivating (“incentive salience” [28]), we performed a conditioned reinforcement probe. In this probe test, rats could press a lever to obtain the CS^+^ or CS^-^ light cue without delivering a sucrose pellet. TBI reduced motivation for the CS^+^ at early and chronic time points post-injury (acute: *p* < 0.001, chronic: *p* = 0.05; Fig 1C). Motivation for this cue decreased at the chronic 108 days post-injury time point for both TBI and sham (*p* = 0.003). Both groups showed low preference for the CS^-^ cue.

To dissociate sign-tracking experience with the lever (Fig 1B) from reduced motivation for reinforcer-predictive cues, we tested a second cohort of rats on an auditory Pavlovian conditioning task at the same acute time point. Auditory cues (Fig 1D) were used without levers or light cues. TBI rats started with lower response rates to the hopper before reaching sham levels (*p* = 0.001; Fig 1E). TBI rats also took longer to discriminate the CS^+^ from CS^-^ (*p* = 0.012; Fig S3A). They were then presented with a conditioned reinforcement probe where they could press the lever to obtain the CS^+^ or CS^-^. TBI rats pressed less for both the CS^+^ and CS^-^ cue (*p* = 0.002, *p* = 0.005; Fig 1F). Overall, these findings suggest that TBI specifically reduces motivation for reinforcer-predicting cues which could represent an inability to integrate predictor cues with their consequences.

### TBI impaired decision-making on a cued gambling-like task

In this experiment, we sought to determine if the altered response to reinforcer-predicting cues observed in Pavlovian conditioning (Fig 1A-F) would also affect operant decision-making in which cues were paired with choice outcomes. We tested rats with TBI on a cued version of the Rodent Gambling Task (cRGT; [29]). We used the same cohort of rats that performed the visual Pavlovian conditioning so that we could correlate performances between the two tests. The cRGT presented 4 choice options via noseports in an operant chamber. Each option had a distinct reinforcement profile with differing probabilities and magnitudes of sucrose pellet “wins” and time-out “losses” (Fig 1G). The four options are a suboptimal less-risky choice (1-pellet; P1), an optimal but low reinforcer choice (2-pellets; P2) and two risky but large reinforcer value choices (3- and 4-pellets; P3, P4). Previously, in a non-cued version of the task, TBI decreased optimal preference but resulted in high variability in resulting preference with some preferring risky choices and some preferring suboptimal [13]. In the cued version used here, the complexity and variability of audiovisual cues increased with reinforcer magnitude (Fig 1G). Prior studies demonstrated that audiovisual cues biased rats toward riskier options [29–31].

Preference for risky versus safer options can be simplified in a single Choice Score variable that subtracts risky from optimal choices. TBI significantly decreased the Choice Score (*p* = 0.001) and attenuated development of safe preference over weeks of testing (*p* = 0.002; Fig 1H). To further explore the specifics of this effect, interdependent choice options were broken down in a Bayesian regression as previously described [32]. This effect was driven by decreased preference of the optimal 2-pellet choice in favor of both the suboptimal 1-pellet and risky 3-pellet options (CI’s not containing 0; Fig 1I). The preference for both suboptimal, minimally-cued, safe 1-pellet option and highly-cued, risky 3-pellet option suggest that the cues are not the main driver of the TBI effect.

To test persistent motivation for the cues paired with winning outcomes on the cRGT, rats were tested for 3 sessions in extinction. During extinction, there was a 100% probability of delivering the cued outcome and no punishment time outs, however, no pellets were delivered. There was no effect of TBI on choice during extinction and choice preference remained stable (Fig S4G-I).

To evaluate relations between Pavlovian behaviors and cRGT decision-making, we performed a series of regressions to compare performance. TBI did not significantly moderate these relationships (see Supplement 2). However, sign/goal-tracking ratios in the acute phase predicted chronic sign/goal-tracking ratio (*p* < 0.001; Fig S6G), and sign/goal-tracking ratio predicted CS^+^ presses at both acute and chronic time points (*p’s* < 0.003; Fig S6I-J). In contrast, cRGT performance was not directly related to Pavlovian behaviors (*p*’s > 0.212; Fig S6A-F).

Together, these data indicate that while TBI impairs behavior in both Pavlovian and operant decision-making contexts, they are dissociable. However, there may be common neural mechanisms leading to both effects.

### TBI reduced markers of behavior-related neuronal activity in the nucleus accumbens core

To better understand which brain regions were impaired after brain injury and contributed to the altered Pavlovian conditioning and decision-making, we chose to broadly screen for activity in key brain regions. We used immunohistochemistry to quantify the transcription factor ΔFosB. ΔFosB expression increases during prolonged neuronal activity [33, 34], making it a useful initial screening tool for behavior-dependent activity. We chose four locations involved in decision-making and motivation for reinforcers: the lateral and medial orbitofrontal cortex for their involvement in decisions regarding probability [35, 36], and the nucleus accumbens core and shell for their role in motivation and reinforcer processing (Fig 2A) [37].

**Figure 2.**
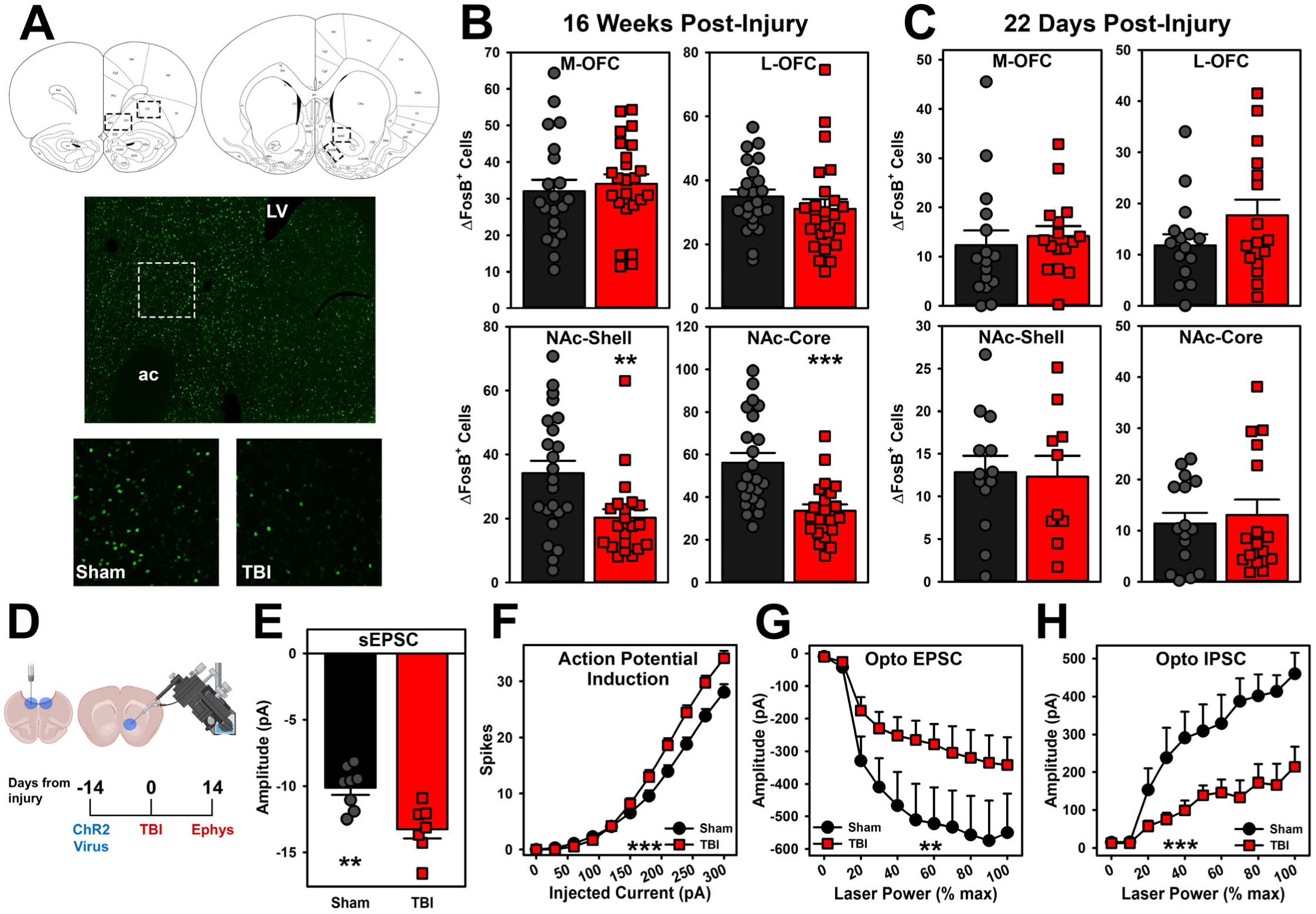
TBI reduced markers of behavior-related neuronal activity, reduced PFC input, and increased neuronal excitability in the nucleus accumbens core. Rats in the behavioral studies (see Fig 1) were evaluated for expression of ΔFosB (panels A-C). A separate cohort received TBI or sham procedures (N = 30) and then at 14 days post-injury were evaluated using slice electrophysiology (panels D-H). A subset of rats also received optogenetic virus for assessing optogenetic-evoked currents (n = 20). A) Atlas showing selected regions-of-interest (ROI) and representative images from NAc core with ΔFosB labeling. B) Quantification of ΔFosB-positive cells after long-term behavior testing (see Fig 1A). TBI reduced expression of the neural activity marker ΔFosB selectively in the NAc core (*p* < 0.001) and shell (*p* = 0.016). C) Quantification of ΔFosB-positive cells after shorter-term (see Fig 1D). TBI did not cause any differences in ΔFosB expression after Pavlovian conditioning. D) Experimental timeline for electrophysiology experiment and depiction of recordings from PFC projections after injury. E) Magnitude of miniature spontaneous excitatory postsynaptic current (EPSC) amplitudes. TBI increased miniature EPSCs (*p* = 0.004). F) Response to current injection. TBI increased neuronal excitability at higher levels (*p* < 0.001). G) Optogenetically-evoked EPSCs from PFC terminal stimulation. TBI decreased the amplitude EPSCs (*p* < 0.001). H) Optogenetically-evoked inhibitory post-synaptic currents (IPSCs) from PFC terminal stimulation. TBI decreased the amplitude of IPSCs (*p* = 0.002). Data are means + SEM with individual subject data superimposed as points on top of bars; *** = *p* < 0.001, ** = *p* < 0.01. Panel A adapted from Paxinos and Watson, 2017, The Rat Brain in Stereotaxic Coordinates, with permission from Elsevier.

In the cohort that performed visual Pavlovian conditioning and the cRGT, there were no differences in the OFC regions (*p*’s > 0.05; Fig 2B), however there was a significant reduction in active cells in the NAc-core (*p* < 0.001; Fig 2B) and NAc-shell (*p* = 0.016; Fig 2B). Because ΔFosB is expressed on a continuum, we increased the minimum brightness threshold to further isolate the “most active” cells. The TBI effect robustly held in the NAc-core (*p* = 0.003 at highest threshold; Fig S7D), but not the NAc-shell (*p* = 0.10 at highest threshold; Fig S7C). Interestingly, the lateral OFC also showed a significant reduction when filtered to the most active cells (*p* = 0.012; Fig S7B). These data suggest that the NAc-core may be a key nexus point for not only the cue-motivation deficits, but also impaired decision-making.

In the cohort that performed just the auditory Pavlovian conditioning task, we performed the same ΔFosB analysis to see if these effects could be observed early and with minimal behavioral experience (only 10 sessions). There was no effect of TBI on ΔFosB expression without the long-term behavioral experience (*p*’s > 0.118; Fig 2C), suggesting that the above data were related to task type or amount of testing.

### TBI reduced PFC input to the NAc core and increased basal and evoked excitation

To better evaluate how the NAc core changes in response to frontal TBI, we performed a subacute electrophysiology experiment. Rats were first injected with an excitatory (ChR2) optogenetic virus in the PFC 14 days prior to their injury, given a TBI (or sham control), then euthanized at 14 days post-injury (Fig 2D). Recordings from NAc slices indicated that TBI increased the amplitude of spontaneous mini EPSCs (*p* = 0.004; Fig 2E), suggesting an enhanced strength of existing excitatory synapses. We also observed an exaggerated response to stimulation of neurons such that more action potentials were evoked with a similar level of stimulation (*p* < 0.001; Fig 2G). Notably, these effects occurred despite other basal membrane properties such as the resting potential and membrane resistance being intact (*p*’s > 0.268; Fig S8). These results suggest that TBI enhanced excitability of neurons within the NAc core.

To determine how much of this change in excitation might be due to loss of PFC input, we optogenetically stimulated the NAc PFC terminals and found that this pathway was no longer able to drive a strong response after TBI (EPSCs: *p* < 0.001; IPSCs: *p* = 0.002; Fig 2G-H).

These data suggest that the increased excitatory drive may be a compensatory response to the loss of input from the PFC. While it is not clear whether this is due to change in projection input strength or local interneuron dysregulation, it could affect integration of inputs from the many regions that synapse onto the NAc.

### TBI reduced transcriptionally-inferred intercellular communication and increased inflammatory, cell stress, and plasticity pathways in the NAc

To assess how loss of PFC input remodeled the NAc, we performed single nucleus RNA sequencing of the NAc in another set of rats with TBI or sham procedures at 14 days post-injury. Rats’ nuclei were barcoded at the subject level, allowing statistical evaluation of variance. After filtering, there were 101,398 neurons for analysis with an average of 3,415 genes/cell identified and 13,428 transcripts/cell across 19 subjects (9 TBI).

To further examine the effects at the level of neuronal subtypes, the dataset was clustered using Seurat. Clustering of individual cells based on canonical markers revealed 23 distinct clusters. Transcriptionally similar ones were combined to yield 1 glutamatergic cluster, 1 cholinergic cluster, 5 MSN subtypes, and 6 other interneuron subtypes (Fig 3A-B). Non-MSN interneurons were then combined for analyses. There was no difference in the proportion of cell types between sham and TBI (*p*’s > 0.099; Fig 3C).

**Figure 3.**
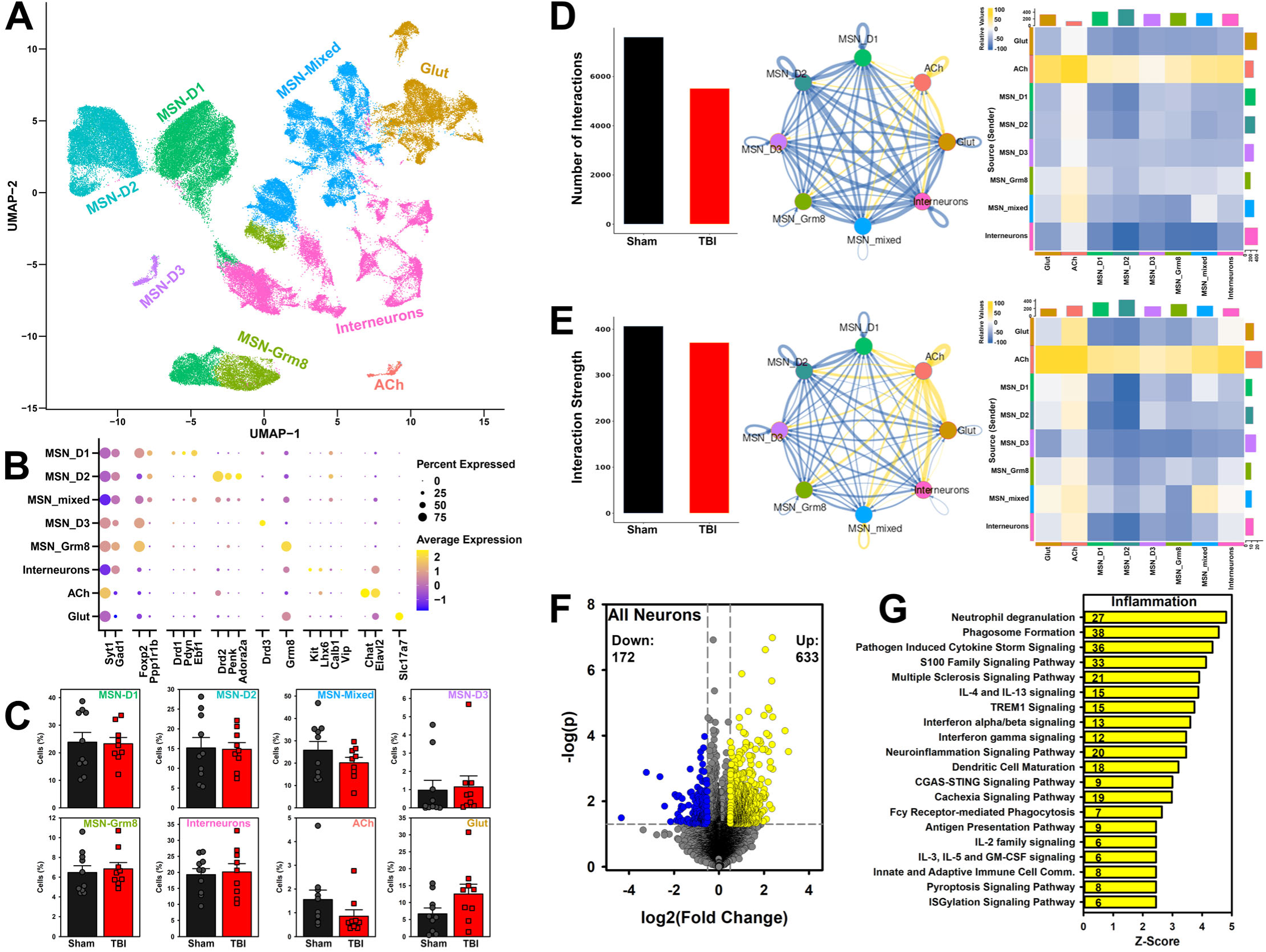
Single nucleus RNA sequencing of NAc neurons at 14 days post-injury. Rats received sham or TBI procedures (N = 19). A) UMAP plot of neurons from NAc displaying clusters of 8 major neuron types. B) Dotplot indicating expression of markers for each neuron cluster (yellow = increased). C) TBI did not alter the proportion of any neuron type (*p*’s > 0.099; means + SEM + subject dots). D-E) Aggregate communication probability inferred from CellChat. Total number of interactions, (D; bar graphs) interaction strength (E) as well as interactions across cell types (chord diagrams, heatmaps; yellow = increased). TBI reduced the total number of inferred interactions (Sham: 7,604; TBI: 5,523) and overall interaction strength (Sham: 406.81; TBI: 370.87). However, the cholinergic cluster had higher outgoing signaling relative to other cells (chord diagrams, heatmaps; yellow = increased). F) Differentially expressed genes across all cells analyzed with conservative pseudobulk approach. TBI increased 633 genes (yellow) and decreased 172 genes (blue). G) Ingenuity Pathway Analysis mapping from differential genes with number of contributing genes superimposed on top of bars. TBI increased inflammatory pathways (43 total, only the top 20 are shown). MSN – medium spiny neurons; ACh – cholinergic neurons; Glut – glutamatergic neurons.

First, we sought to understand the global effect of TBI on the NAc transcriptome. CellChat [38] was used to infer cell–cell communication networks by mapping putative ligand–receptor interactions based on gene expression profiles. TBI reduced the total number of inferred interactions across the network (Sham: 7,604; TBI: 5,523), suggesting a global decrease in neuronal intercellular communication that was broadly consistent across most cell types. In contrast, cholinergic interneurons exhibited a distinct pattern, with increased outgoing signaling to multiple cell types relative to other populations (Fig 3D), suggesting a potential compensatory or regulatory role following brain injury. The overall interaction strength of interactions was less affected by TBI (Sham: 406.81; TBI: 370.87), but a similar relative increase in cholinergic signaling was observed in this metric as well (Fig 3E).

To evaluate global gene expression changes and provide a broad picture to complement the CellChat data, pseudobulk analyses were performed to mitigate power inflation given the large number of individual neurons. Transcripts were summed at the subject level and then pseudobulk differential gene expression was performed (minimum change *z* = 0.5). A total of 805 genes were differentially regulated by TBI (633 increased, 172 decreased; Fig 3F; Supplement 2). These differential genes were used in Ingenuity Pathway Analysis to evaluate pathway enrichment. The pathway list was filtered to remove non-neural paths and remove any with a z score below 0.5. 72 total pathways were affected (59 after B-H correction). Of these, 43 were directly related to inflammatory responses and all were increased (Fig 3G). The remainder included pathways related to cell stress, extracellular matrix reorganization, and intra/extra-cellular signaling (Supplement 2). Collectively, these data indicate an RNA profile of inflammatory stress and consequent plastic reorganization at a subacute post-injury time point.

### TBI strongly affected individual neuron subtypes in the NAc

To make use of the high-resolution cell-type data, we performed differential gene expression (DEG) analysis for each neuronal cluster. All DEGs were analyzed using pseudobulk approaches to control false positives. All cell types showed differential regulation of genes due to TBI with particularly pronounced effects in the D1, D2, and mixed MSNs (Fig 4A). Cell types not discussed here are presented in Supplement 1 (Fig S9). We then determined which DEGs were commonly affected across all cells and particularly the key MSN subtypes.

**Figure 4.**
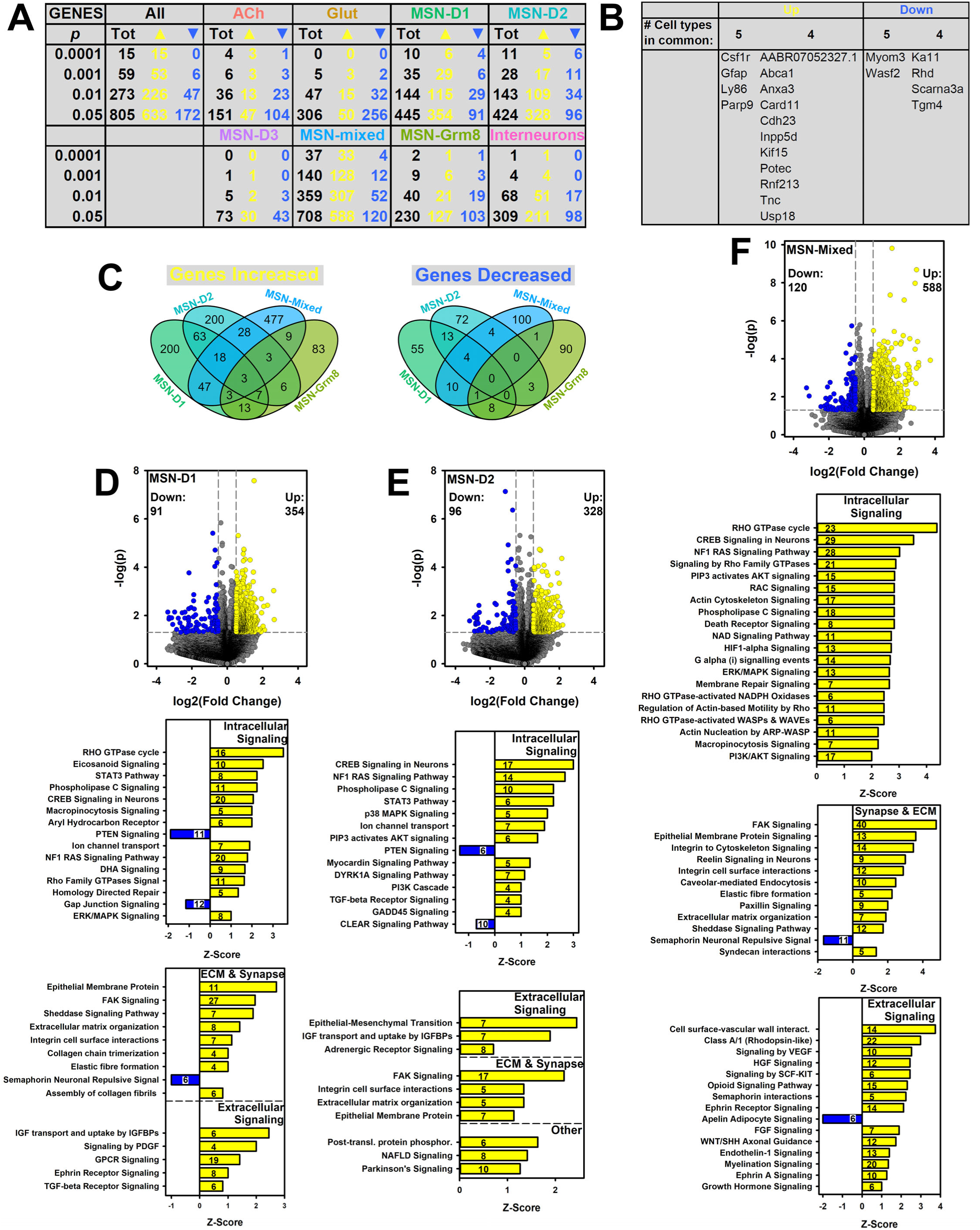
Pseudobulk analysis of differentially expressed genes (DEGs) and corresponding pathways across neuron types broken out from data shown in Figure 3. **A)** TBI affected all neuron types, even when stricter significance cutoffs were used. B) TBI increased 15 genes and decreased 6 genes across 4 or more cell types. C) Venn diagram showing unique and overlapping increased or decreased genes in MSNs. D-F) TBI changed gene expression for MSN-D1, MSN-D2, and MSN-mixed cells. These mapped largely onto plasticity-related pathways, including intracellular signaling, matrix modification, synapse modulation, and other types of extracellular signaling. The number of contributing genes is superimposed on top of bars. MSN – medium spiny neurons; ACh – cholinergic neurons; Glut – glutamatergic neurons.

Overall, individual neuron subtypes were strongly influenced by TBI. A total of 345 genes were either increased or decreased in more than one cell type. The most common DEGs are shown in Fig 4B, with several appearing in 4 to 5 cell types. The common increased DEGs were largely inflammatory but also included plasticity-related genes. Surprisingly, TBI increased canonical microglial and astrocytic genes *Csf1r* and *Gfap* in these putative neurons. Examination of expression revealed low mean normalized counts relative to true microglia or astrocytes (*Csf1r*: 0.31 in neuron vs. 15.47 in microglia; *Gfap*: 0.48 in neuron versus 5.88 in astrocytes). This suggests some minor ambient RNA contamination, which can be more pronounced in injury conditions [39] and may reflect aspects of injury vulnerability. Overall, the MSNs had several common DEGs affected by TBI, however, there were many more genes uniquely affected in specific cell types (Fig 4C).

### TBI drove large effects in canonical D1- and D2-medium spiny neurons

TBI substantially affected gene expression in MSN-D1 and MSN-D2 cells (Fig 4A, D, E) mapping to 45 and 27 affected pathways, respectively. Notably, these cells had less inflammatory pathways (MSN-D1: 11; MSN-D2: 4). Instead, most pathways indicated increased intracellular signaling (e.g., PLC and CREB signaling), extracellular matrix modification, and other extracellular signals (e.g., neurotransmission). Notably, these pathways were almost entirely increased, suggesting cells were undergoing a large-scale plastic change at this time point. Full pathway details and summaries are described in Supplement 2.

### TBI most strongly affected an indistinct type of medium spiny neuron in the NAc

One of the strongest effects on gene regulation was in a cluster of MSNs with an indistinct phenotype that included D1 and D2 markers (“MSN-mixed”; 708 DEGs: 588 up, 120 down; Fig 4F). These cells had one of the largest inflammatory responses (57 increased pathways; Supplement 2), likely driving the global inflammatory effect (Fig 3G). Despite this, most of the transcriptional changes were spread across a variety of pathways (79 additional enriched pathways Fig 4F). These included 31 intracellular signaling pathways, 15 extracellular signaling pathways, and 12 synaptic and extracellular matrix pathways. Full pathway details and summaries can be found in Supplement 2. Overall, these findings suggest a particularly vulnerable cell type with high inflammatory/stress response to brain injury and high plasticity.

This mixed phenotype is unusual and not well represented in pure NAc core samples. It is likely that our larger punch also captured NAc shell and that is what is represented here [40]. But given the large differential response to TBI, it could also indicate a cluster of cells driven by injury effects that were common to both D1 and D2 MSNs. To evaluate this, we plotted several genes identified in D1 and D2 MSN subclusters from recent rat single nucleus RNA sequencing datasets: *Myo3a, Fstl4, Dach1, Scube1, Stk32a, Htr4, Ebf1, Ppm1e, Reln, Htr7, Wls, Reln* [41, 42]. None of these distinguished this mixed cluster as more D1- or D2-like; it had gene expression for several overlapping D1, D2, and even Grm8-MSNs (Fig S10).

### Accumbens glutamatergic and cholinergic cells showed an opposing effect in response to TBI

Cholinergic interneurons represented 0.74% of the neurons and glutamatergic neurons represented a surprising 11.65%. The larger-than-anticipated glutamatergic population may represent a punch that encompassed regions slightly beyond the NAc core. Both glutamatergic and cholinergic neurons had more decreased genes than MSNs (Fig 4A; Fig S9A-B). This translated to most mapped pathways being decreased as well (Fig S9A-B). Full pathway details and summaries can be found in Supplement 2. These interneurons may be providing some level of compensation for the major changes seen in the MSNs. The cholinergic changes also corresponded to the increased intercellular communication in these cells post TBI (Fig 3D-E).

### TBI blunted NAc calcium activity in response to reinforcement-predictive cues

In the final experiment, we tested whether NAc activity was reduced during relevant periods of behavior due to TBI. The data above suggested substantial subacute reorganization (Fig 3-4) and altered excitability (Fig 2E-F), however the conclusion from our behavioral screen was that the NAc was underactive in behaving rats with TBI (Fig 2B). To address this, we performed fiber photometry using a calcium reporter (GCaMP6f) in the NAc core after TBI or sham procedures (Fig 5A) to obtain a real-time readout during behavior. Rats received virus 14 days prior to photometry probe implant and TBI procedures. After 21 days of recovery, they underwent visual Pavlovian conditioned approach (Fig 1A) while recording calcium signal. The task was slightly modified from above (from 5 s to 8 s cue) to obtain more recording signal.

**Figure 5.**
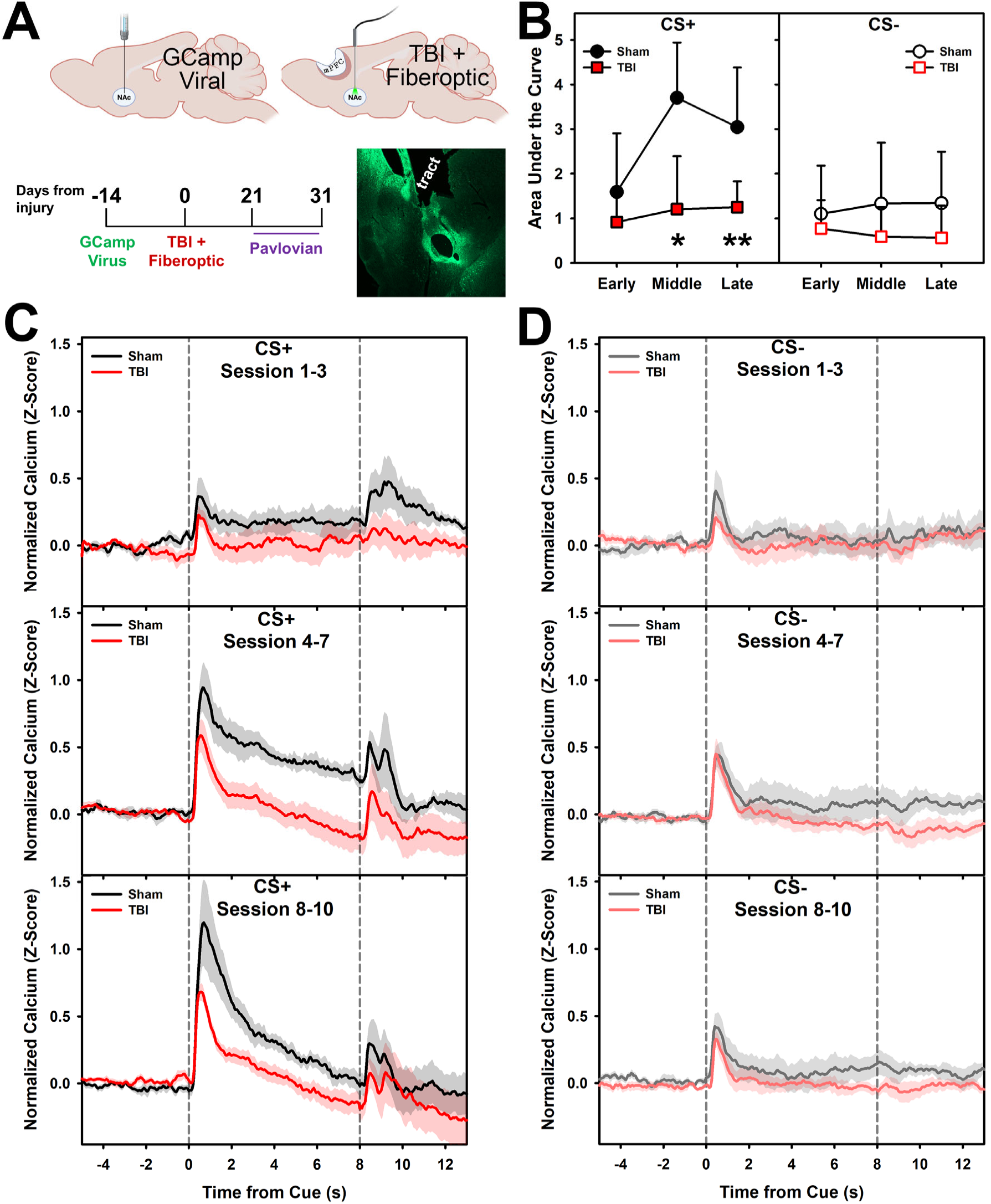
TBI reduced NAc calcium responses to reward-predictive cues. Rats were given a GCaMP virus, then TBI or sham procedures and evaluated on visual Pavlovian conditioning (N = 12). A) Experimental timeline, depiction of manipulations, and representative viral image with tract next to lateral ventricle. B) Quantification of area under the calcium response curve for CS+ and CS− cues across early (session 1-3), middle (session 4-7), and late (session 8-10) phases. Sham rats developed larger CS+ responses over training, whereas TBI rats showed attenuated responses (*p*’s < 0.037). Responses to the CS− cue were similar between groups. C) Average calcium traces aligned to CS+ cue onset (first dashed line) and reward delivery (second dashed line) across learning phases. D) Average calcium traces aligned to CS− cue onset and where reward delivery would have occurred during a CS+ for reference. Data are means + SEM (error bar or error shading); ** = *p* < 0.01, * = *p* < 0.005.

In the early phase of the task (sessions 1-3), there were no differences between signals in sham or TBI rats for the CS^+^ cue (*p* = 0.341; Fig 5B-C). In the middle phase of learning (sessions 4-7) shams began to show a larger response to the CS^+^ cue as measured by the area under the curve and this was attenuated by TBI (*p* = 0.007; Fig 5B-C). This difference persisted into the late learning phase (sessions 8-10; (*p* = 0.037; Fig 5B-C). There were no differences in peak values, though this may be due to higher variance in sham peaks (*p* = 0.084; Fig 5C, S11A). Both groups showed a similar response to the CS^-^ and discriminated between the two (*p*’s > 0.183; Fig 5D). There were no differences in the latency to peak value (*p* = 0.987; Fig S11B).

However, the peak for TBI rats decayed back to baseline faster (*p* = 0.021; Fig S11C). Together, these data indicate an underactive NAc in relation to processing signals predicting reinforcement, but not reinforcement itself. This impairment may contribute to lower motivation for reinforcer-predictive cues (Fig 1C, F) but may also impair learning about choice outcomes, providing a potential explanation for the poor behavioral performance in the cRGT (Fig 1H).

## Discussion

Traumatic brain injuries are a key risk factor for psychiatric and neurological diseases. In the current study we used one of the simplest reinforcement-driven behaviors, Pavlovian conditioning, to isolate a fundamental symptom associated with TBI: an altered response to cues predicting reinforcers (Fig 1A-F) [17]. Lost or dysfunctional inputs from the damaged mPFC (Fig 2G-H) significantly altered the transcriptional landscape of the NAc (Fig 3-4), ultimately leading to reduced activity in anticipation of reinforcers (Fig 5B-C). This may indicate a reduced salience of environmental events which could ultimately contribute to TBI-driven impairments in decision-making (Fig 1G-I), behavioral flexibility, and learning. However, if the NAc is a key nexus point for these impairments, it also opens opportunities for targeted treatments.

Given the innervation of the NAc core from the mPFC [43], it is not particularly surprising that mPFC damage would cause a loss of input. However, the functional consequences of this loss were not well understood. Despite being distal to the injury, the NAc showed a robust inflammatory response to TBI (Fig 3G). This is consistent with progressive secondary pathology [1, 44] and potentially driven by dying and degenerating projections. However, the transcriptional profile indicated more than mere inflammation with high upregulation of intra- and extra-cellular signaling and synaptic pathways (Fig 4). This was paired with functional changes in physiology, including compensatory increased basal excitability and response to stimulation (Fig 2E-F). The PFC normally has a net inhibitory effect on the NAc via interneuron activation [45]. Thus, the increased excitation observed here could indicate an early signature of NAc dysfunction where it loses its ability to efficiently integrate information across input sources. Losing the ability to parse the diverse inputs to the NAc could generate a multitude of behavioral impairments, including impaired decision-making, impulsivity, and even risk for substance use disorders [20, 46, 47]. Indeed, we have observed several of these in our model of injury – rats with frontal TBI are more impulsive and are impaired in multiple aspects of decision-making and behavioral flexibility [12–15].

A classic view of the NAc postulated it as the “reward center” [48]. This was further refined as dopaminergic inputs were identified to be robust prediction signals for reinforcers [49]. Later studies emphasized its role parsing both appetitive and aversive stimuli. These suggested that the difference between D1 and D2 MSNs in the NAc may be in how they encode outcome valence, differing in their roles projecting along the direct and indirect pathways, respectively [50]. However, recent work challenges the simplicity of a D1/D2 opposition model. For example, mice preferred optogenetic self-stimulation for brief, but not prolonged stimulation, regardless of whether it was D1 or D2 MSN stimulation [51]. Further, both MSN types responded to cues signaling an operant response regardless of reinforcer valence (sucrose vs. shock avoidance) [52]. Given the wide-scale behavioral dysfunction after TBI, the role of the NAc as a locus for processing salience of outcomes makes it an attractive target for developing potential therapeutics. In the current study, TBI attenuated the NAc response to predictive cues (Fig 5B-C), but more work will be needed to determine cell-type specific contributions.

In the current transcriptomic data, TBI altered expression of hundreds of genes across most major NAc neuron subtypes (Fig 4A). TBI did not change absolute proportions of cells, suggesting there was no “die-off” off due to reduced PFC input (Fig 2G-H). However, CellChat analysis indicated fewer inferred interactions across the network suggesting an overall suppression of intercellular communication after TBI. When this was broken out by cell type, cholinergic interactions showed an opposing effect with (relative) increased interactions (Fig 3D-E). This oppositional effect of cholinergic cells was also reflected in gene expression, with more suppression of genes and pathway (Fig S9A) relative to other subtypes. When all neurons were considered together, some of the most dominant gene expression changes were in pathways linked to neuroinflammation (Fig 3G). Transcriptomic changes were most pronounced in D1 and D2 MSNs, but also a mixed MSN phenotype which had both D1 and D2 markers (Fig 4D-F).

Mapping of known D1 and D2 markers did not help to resolve the identity of these cells (Fig S10); they remained mixed. TBI elevated inflammatory pathways in each of these MSNs, but also increased plasticity-associated pathways (intracellular signaling, extracellular signaling, synaptic change, extracellular matrix) suggesting some level of reorganization due to these stresses. This was most pronounced in the MSN-mixed phenotype, suggesting they may represent a particularly vulnerable population of MSNs. Though this mixed MSN phenotype has not been widely reported, it may be due to NAc shell tissue in the sample [40] which is likely due to the size of the glutamatergic population (Fig 3C).

To date, no experiments have examined the transcriptome of the NAc in the context of brain injury. The bulk of research on the NAc transcriptome is in substance abuse and related literature. However, TBI also alters the response to drugs of abuse such as cocaine. TBI accelerated cocaine self-administration acquisition [53] and enhanced cocaine conditioned place preference [54, 55]. In uninjured animals, non-contingent cocaine altered gene expression, particularly in D1 MSNs [56], including a “responsive” subcluster enriched for the reelin gene *Reln* [57], a regulator of synaptic plasticity [58]. Although our dataset shows increased extracellular matrix and plasticity pathways (Fig 4D-F), consistent with heightened vulnerability, *Reln* was decreased across all MSN subtypes after TBI (Fig S12; Supplement 1). In contrast to the effects of non-contingent cocaine, operant training for food reinforcers drove gene changes that were particularly pronounced in D2 MSNs and which mapped to many synaptic and intracellular plasticity-related pathways [59]. Brain injuries represent a particular challenge to integrate with fundamental research as the changes are widespread across many subtypes of cells. However, the large-scale shifts across MSNs in the current data (Fig 3-4) have the potential to set up both adaptive and maladaptive responses to brain injury.

An interesting nuance of the current behavioral data is that while TBI attenuated NAc signal to the reinforcer-predicting cue (Fig 5B-C), it did not merely reduce behavioral output. Instead, we observed a strong shift toward goal-tracking behaviors (Fig 1B). This behavior, while aberrant, is not inherently maladaptive but reflects reduced motivational properties of these cues (Fig 1C, F). When tested on the cRGT, rats were required to integrate information about probability, magnitudes, and cued outcomes and adjust their behavior accordingly. TBI reduced the ability to optimize this choice process (Fig 1H-I), however the cues did not exert any clear influence. TBI increased choice of both the simplest cue (P1, suboptimal) as well as a complex cue option (P3, risky) but did not increase preference for the highest cue complexity option (P4, risky). Ultimately, there was no clear relation between sign- or goal-tracking and cRGT choice behavior (Fig S6) despite robust injury differences in both. This contrasts with a prior study which indicated a modest relationship for these variables [60]. Together, these behavioral data suggest that while rats with frontal TBI can learn simple contingencies, there is aberrant processing in the NAc leading to difficulty adapting behavior to outcomes, particularly when those outcomes are more ambiguous as in the probabilistic cRGT.

The findings in the current study highlight that TBI-induced dysfunction in the NAc arises not simply from loss of input, but from a functional reorganization of the structure around this change in input. This may ultimately alter how information is integrated across cell types, circuits, and signaling systems. The substantial remodeling of the NAc poses many interesting questions to better understand the recovery process. Given the large-scale behavioral deficits linked to poor NAc function (Fig 1), it is tempting to assume that the early changes are maladaptive. In particular, the early inflammatory response (Fig 3G) could contribute to long-term functional deficits and represent a potential treatment target. However, it is crucial to also consider the highly-plastic cellular environment as this region adapts to a sharp reduction in PFC input. Thus, future studies will need to determine how specific neuronal populations, afferent pathways, and cell-level adaptations contribute to this and which might instead be harnessed to treat dysfunction. Specifically, it may be possible to restore some NAc function through targeted receptor modification, direct stimulation, or indirect stimulation of specific pathways (e.g., mesolimbic dopamine pathway). Indeed, one study found that deep brain stimulation of the NAc after TBI improved learning in the Morris Water maze [51]. However, it remains to be seen whether this is sufficient for situations with more subtle outcome cues such as the probability-based decision-making measured here (Fig 1G-I). Overall, the current findings establish the NAc as a key nexus for behavioral deficits resulting from frontal TBI and open many potential avenues to develop effective therapeutics.

## Methods

### Research Design

The following experiments evaluated the role of NAc dysfunction after frontal TBI. Experiment 1 – Effects of TBI on Visual Pavlovian Conditioning, Conditioned Reinforcement, and Cued Rodent Gambling Task (Fig 1A, G). This experiment served to establish the key behavioral deficits relating to processing of cues and environmental outcomes after TBI and used histology to identify the NAc as a major potential contributor. Experiment 2 – Effects of TBI on Auditory Pavlovian Conditioning and Conditioned Reinforcement (Fig 1D). This experiment confirmed that deficits in motivation for a conditioned cue were not just due to spatial orientation in the prior experiment. Experiment 3 – Effects of TBI on Nucleus Accumbens Electrophysiological Properties and PFC Input (Fig 2D). This experiment evaluated loss of projections from the PFC and consequences on cellular properties. Experiment 4 – Effects of TBI on Nucleus Accumbens Cell-Level Neuronal Transcriptome. This experiment evaluated how individual cells in the distal accumbens were remodeled by brain injury. Experiment 5 – Effects of TBI on Task-Related Nucleus Accumbens Activity (Fig 5A). This experiment linked the observed functional deficits to a real-time readout during behavior to confirm that the NAc was not processing information in the same fashion after injury.

### Subjects

Subjects were 140 Long-Evans rats (65 female; Charles River, Wilmington, MA) after 5 exclusions for poor viral expression. Rats were approximately 3-4 months old at the time of TBI. Rats were pair-housed in standard Allentown cages (Allentown, NJ) and single housed after injury. Rats were housed in an AAALAC accredited vivarium on a 12 h light/dark reverse cycle with behavioral rats food-restricted to 85% of free feeding and non-behavioral rats fed ad libitum. All had free access to water. All work was approved by Ohio State University Institutional Animal Care and Use Committee (IACUC protocol: 2021A00000037).

### Viral Infusion Surgery

Rats were anesthetized with isoflurane at 5% in 1-2 L/min oxygen. Rats were placed on a stereotaxic frame and maintained at 2-4% of isoflurane in 200-400 mL/min air. General analgesic (carprofen, 5 mg/kg) and local analgesic (bupivacaine, 0.1ml at 0.25% concentration) were administered subcutaneously. The incision site was sterilized, and a midline incision was performed with a scalpel, and retracted. After measuring from bregma, a burr hole was made above the target location and a 10 µl Nanofil Syringe (World Precision Instruments, Sarasota, FL) attached to an electronic pump (UPM3, World Precision Instruments) used to lower to the target.

For optogenetic experiments, the target was the bilateral prelimbic cortex (AP/ML/DV: +3.2, ±0.8, −3.4 mm from bregma) and 0.5 µl of virus (pAAV-hSyn-hChR2(H134R)-mCherry in AAV9, titer >= 1*10^13^, Addgene #26976, Watertown, MA) was infused at 100 nL/min with an additional 5 minutes at the end for diffusion. For calcium imaging photometry experiments, the target was the nucleus accumbens core (AP/ML/DV: +1.6, +1.6, −7.2 mm from bregma) and 0.8 uL of virus (pENN.AAV.CamKII.GCaMP6f.WPRE.SV40 in AAV9, titer 3.1*10^13^, Addgene #100834) was infused at 75 nL/min with an additional 8 minutes at the end for diffusion.

After surgery, the surgical site was sutured. Triple antibiotic ointment was applied to the incision site. Carprofen was given 24 h after brain injury and rats monitored for recovery. TBI surgeries occurred 2 weeks after viral surgeries.

### Controlled Cortical Impact (TBI) Surgery

Rats were randomly assigned to receive a TBI (n = 71) or sham surgery (n = 69) with roughly equal sexes in each group except for the photometry experiment (all male). TBI rats received a moderate to severe bilateral injury to the mPFC via controlled cortical impact [61]. Rats were prepared as described above (analgesia, placed in stereotax), and then a 6 mm diameter craniectomy was performed at +3 mm from bregma, centered on the midline. An impactor (Neuroscience Tools, O’Fallon, MO) and driver (Leica Biosystems, Buffalo Grove, IL) were used to induce a bilateral TBI at 3 m/s, at a depth of −2.5 mm and 500 ms dwell time. Bleeding was stopped and triple antibiotic ointment was applied on the incision site. Sham surgeries were identical to TBI but excluded the craniectomy and impact.

### Fiberoptic and Combined TBI/Fiberoptic Implant Surgery

Additional protocol details (including troubleshooting for implants on top of a large craniectomy) are available in Supplement 1. In brief, rats were given a CCI as described above. Prior to craniectomy, a new reference point was marked on the skull (−3 mm from bregma). Four to five screws were placed in front of and behind the craniectomy. A fiberoptic probe (10 mm length, 400 μm diameter, Thor Labs, Newton, NJ) was lowered to 0.2 mm above the viral location (AP/ML/DV: +1.6, +1.6, −7.0 mm from bregma). The craniectomy was then covered and sealed with a UV-cured dental cement (Natural Elegance Premium Flow Composite, Henry Schein, Melville, NY). C&B Metabond (Parkell, Edgewood, NY) was then used to form a base layer for the head cap on top of the craniectomy, against the exposed fiberoptic, and under the screws. After curing, the remainder of the headcap was built with standard dental cement (Jet Powder/Liquid, Lang Dental, Wheeling, IL). Skin was loosely sutured around the base of the headcap to maintain access. Sham procedures did not use a craniectomy to help mitigate headcap loss.

### Behavior Apparatus

Operant behavior testing took place in Med-Associates chambers (St. Albans, VT). These contained two levers with a light above each and pellet hopper in the middle. Some chambers also contained an array of 5-choice holes on the wall opposite the levers. Sucrose pellets were used as reinforcers (Bio-Serv, F0021).

### Visual Pavlovian Conditioning Behavior

Rats were put through a conditioned approach paradigm similar to previous descriptions (Fig 1A) [17, 62]. For a given trial, either the left or right lever would extend into the chamber, accompanied by the illumination of the cue light above the respective lever. After 5 s (8 s in the photometry experiment), the lever retracted, and the light turned off. One lever (counterbalanced between subjects) was associated with a single reinforcer delivery and illumination of the tray light immediately after (designated CS^+^), while the other was never paired with reinforcement (designated CS^-^). Lever presentation was pseudorandomly determined such that rats received equal exposure to CS^+^ and CS^-^. Trials were presented on a variable time (VT)-60 schedule for one-hour sessions. The primary variables of interest were the total time spent at the lever and hopper, and the ratio between hopper and lever times during CS^+^ presentation. The primary variables of interest were sign- and goal-tracking behaviors which are measured by the time spent on/depressing the lever and the time spent in the hopper, respectively. These were then represented as the (log-transformed) ratio between the time spent in these behaviors to capture a continuous measure of the degree to which a rat engaged in each.

### Auditory Pavlovian Conditioning Behavior

Conditioning procedures were identical to the visual conditioning, however the CS^+^ and CS^-^stimuli were a 1 kHz and 5 kHz tone (counterbalanced between subjects; Fig 1D). No levers were extended at any point. Hopper time was tracked to determine degree of conditioning.

### Conditioned Reinforcement Behavior

After visual or auditory conditioning, rats underwent one Conditioned Reinforcement probe to measure willingness to press a lever on an FR-1 schedule to receive either the CS^+^ or CS^-^. No sucrose pellets were delivered. Rats in the visual conditioning experiments underwent one 30-min session, while rats in the auditory underwent one 180 min session. Lever presses for CS^+^ and CS^-^ were the primary dependent variables.

### Cued Rodent Gambling Task (cRGT)

The cRGT was carried out as previously described [29]. Sessions lasted 30 minutes each. In brief, rats were trained to nose poke in the holes for reinforcers. They were then given 7 sessions of “forced choice” to expose them to the probability, magnitude, cue outcomes of the various cRGT choices. The four choice outcomes had distinct sets of probabilities of “wins” (sucrose pellets) and “losses” (time out from earning) which are depicted in Figure 1G. The audiovisual cues were given during “winning” trials and scaled in complexity and variability with the magnitude of winning outcomes. Besides choice data, many other variables were recorded and are presented in Supplement 1.

### Euthanasia and Tissue Collection

At the conclusion of behavior for histology and immunohistochemistry experiments, rats were transcardially perfused with 0.1 M PBS, followed by 3.7% formaldehyde. The brains were extracted and post-fixed with 3.7 % formaldehyde for 24 h prior to storage in a 30% sucrose + 0.02% sodium azide solution. Brains were frozen and section at 30 μm using a sliding microtome. Slices were then selected for immunohistochemistry or mounted on glass slides for Nissl staining.

For electrophysiology and sequencing experiments requiring fresh tissue, rats were briefly anesthetized using 100% CO₂ introduced into a 24.5-L chamber at a flow rate of 10 L/min (∼41% chamber volume displacement per minute) for at least 1 minute and until unresponsive. Rats were then removed from the chamber, rapidly decapitated, and fresh brains extracted. Brains were either put immediately on ice-cold solution for electrophysiology or coarsely sliced and punched on ice.

### Histology and Immunohistochemistry

Nissl staining was performed with thionin to measure remaining brain volume and validate injuries. Briefly, slides were rehydrated using a series of washes of decreasing ethanol concentrations, placed in thionin solution, then dehydrated with a reverse sequence. Slides were then cover slipped and dried. The slides were digitally scanned (600 DPI) and brain slices surrounding the lesion (+1.0, +2.0, +3.0, +4.0, and + 5.0 mm from bregma) were measured using ImageJ (NIH, Bethesda, MD). Brain volumes were estimated by multiplying the average size of the slices by their thickness (30 um). No injury exclusions were necessary.

ΔFosB immunohistochemistry was performed to quantify neuronal activation using a free-floating fluorescent staining protocol. Briefly, slices were rinsed in dH₂O and incubated in 5% normal goat serum (NGS) in 0.3% Triton PBS for 4–24 h at 4°C under agitation. Slices were then incubated in primary rabbit anti-ΔFosB antibody (Cell Signaling D3S8R; dilution 1:2000 in NGS) for 48 h at 4°C. After rinsing in PBST (0.1% Triton PBS) followed by PBS, slices were incubated in goat anti-rabbit IgG Alexa Fluor 488 secondary antibody (1:2000 in NGS; Abcam ab150077) for ∼12 h at 4 °C in a darkened environment. Following secondary incubation, slices were rinsed in PBST and dH₂O, then mounted using Fluoroshield with DAPI and cover slipped. Regions of interest were imaged at 40x magnification using a Nikon Eclipse Ti confocal microscope and camera. 8 images per region per subject were represented bilaterally and incorporated in the analysis. ΔFosB-positive cells were counted using ImageJ’s automated Analyze Particles function with a minimum area. To get “low”, “medium”, and “high” activity cells, we applied a pixel intensity minimum threshold at three levels which were determined by manual review for each region (range 300–1900).

### Patch-Clamp Slice Electrophysiology

In brief, whole brains were collected at 14-days post-injury and placed into ice-cold sucrose-based cutting solution containing (in mM) 234 sucrose, 2.5 KCl, 1.25 NaH_2_PO_4_, 26 NaHCO_3_, 10 MgSO_4_, 0.5 CaCl_2_, 11 glucose, 1.3 L-sodium ascorbate (PH; 7.3-7.4). Coronal brain sections (300 um) were cut on a Leica VT1200S. Sections with clear NAc were placed into a recovering chamber containing artificial cerebrospinal fluid (aCSF in mM; 124 NaCl, 3 KCl, 2 CaCl_2_, 1 MgSO_4_, 24 NaHCO_3_, 1.25 NaH_2_PO_4_, 10 glucose) at 35°C for 30 min and then moved to room temperature for 40 minutes. Individual hemislices were transferred to a recording chamber perfused with carbogen aCSF and recordings were conducted with patch electrodes (TW150-3, World Precision Instruments) filled with internal solution (140 K-gluconate, 3 KOH, 1 MgCl_2_, 1 CaCl_2_, 5 EGTA, 10 HEPES, 2 Mg-ATP, PH; 7.1-7.3). Cells distributed within NAc core were visualized with infrared differential interference contrast (IR-DIC) optics equipped with (Olympus BX51WI) upright microscope. A MultiClamp 700B amplifier, digidata1550B digitizer, and Clampex11 acquisition software (Molecular Devices) were employed to conduct recording. The signal was filtered at 1 kHz and digitized at 10 kHz.

Whole-cell recordings were achieved by rupturing the membrane under the electrode after obtaining a giga-seal. Cell and electrode capacitances, and serial resistance were compensated automatically. Serial resistance was controlled below 20 mΩ. Voltage-clamp recording was conducted at −70 mV holding potential and delivery of 12 stepwise sweeps (−70 mV to +40 mV) of 400 ms duration with 5 s intervals. Recordings were switched from voltage-clamp to current-clamp recording, and the resting membrane was measured. The input resistance was calculated from 5 evoked voltage response as 200 ms current injections decreased from −100 to −20 pA. The threshold current to evoke the first action potential was determined from a series of 1 s injected current pulses gradually increased in 30 pA increments (11 steps, intervals 10 s). Action potential frequency was measured via injected current curves for each cell. All these recordings were used to evaluate cells excitability. After conducting this current-clamp recording, recording was switched back to voltage-clamp. Spontaneous excitatory postsynaptic currents (sEPSC) were recorded at −70 mV holding potential and spontaneous inhibitory postsynaptic currents (sIPSC) at 0 mV holding potential with each recording lasting 2 min. Clampfit 11 software was used to analyze all data off-line.

For optogenetics testing, a 470 nm light pulse was generated via a dual opto-led power supply (Crian, UK) and 40X water immersion lens to illuminate the recording field. The recording procedure was controlled by patch-clamping software. To measure the absolute oIPSCs (0 mV holding potential) and oEPSCs (−70 mV holding potential), 0.5 mS blue light pulses were delivered in a series of progressively increasing stimulation intensity (0-1 unit) to evaluate the ChR2-positive afferent fibers’ effects.

### Nuclei Extraction and Fixation for Sequencing

After decapitation, brains were coarsely sliced (1 mm slices) in a brain matrix and nucleus accumbens extracted with a 2 mm punch, then snap frozen with dry ice. Frozen tissue was placed into 4 ml of cold enzymatic lysis and nuclei extraction buffer (Miltenyi Biotech, Bergisch Gladbach, Germany) with an RNase inhibitor (0.2 U/µl, ThermoFisher Scientific, Waltham, MA). Tissue digestion was then performed in a GentleMACS dissociator with Octo Cooler (Miltenyi) in a C tube using the program 4C_nuclei_1. Upon completion, tubes were washed with 2 ml lysis buffer and filtered with a 100µm MACs SmartStrainer (Miltenyi) and centrifuged. Nuclei were then resuspended in 1.5ml of resuspension buffer (PBS, 0.04% BSA, 14% nuclei extraction buffer, 0.2 U/µl RNase inhibitor. Nuclei concentration was measured using the LUNA FL florescence cell counter (Logos Biosystems, Gyeonggi-do, South Korea) and the Acridine Orange/Propidium Iodide Stain (Logos Biosystems) as per manufacturer’s instructions. Anti-nucleus microbeads (Miltenyi) was applied to nuclei suspension as per manufacturer’s instructions. Nuclei were then magnetically separated using LS Columns (Miltenyi Biotech) and collected.

A maximum of 1×10^6^ nuclei per sample were fixed using the Parse Bioscience Evercode Nuclei Fixation Kit V3 (Parse Bioscience, Seattle, WA) as per manufacturer’s instructions. Fixed nuclei were resuspended in 100µl of Storage Buffer and passed through a 70µm filter and stored at −80°C until barcoding.

### Single Nucleus RNA Barcoding and Sequencing

Single nucleus RNA sequencing was performed as previously described [63]. Fixed nuclei were barcoded with the Evercode WT Mini V3 and Evercode WT V3 (Parse Bioscience) following manufacturer’s instructions. A total of 15,000 fixed nuclei were barcoded using the WT Mini kit and 120,000 fixed nuclei were barcoded using the WT kits. Equal number of nuclei were barcoded from each sample and upon completion of barcoding, two unique cDNA libraries were generated from the WT Mini kit, while eight unique cDNA libraries from WT kits. cDNA libraries were sequenced at the Michigan Genomics Core (The University of Michigan) or Parse Biosciences for Illumina sequencing at 40,000 reads/cell.

### Single Nucleus RNA Data Processing

After sequencing, FASTQ files were generated from each experiment. The Parse Biosciences pipeline (v 1.5.1) was used to identify transcripts from individual cells, map them to the rat genome (mRatBN7.2), and remove doublets. The resulting count matrices were imported into R and analyzed using Seurat (v.5.3.0). Files were merged into a single object and lack of batch effects confirmed by examining clustering. After examining distributions on violin plots, nuclei with <300 and >9000 expressed genes were excluded to remove debris and doublets, respectively. Nuclei with <300 and >40000 transcripts were also excluded, and due to very low mitochondrial contamination, nuclei with >0.5% of mitochondrial RNA were excluded. In total, 101,398 high quality nuclei were obtained. After filtering, data were normalized (LogNormalize) and scaled, and highly variable genes were computed (vst) and used to compute PCA.

Glial versus neuronal cells were then annotated using a list of markers (Arhgap15, Csf1r, Cx3cr1, Mbp, Mobp, Mog, Enpp6, Pdgfra, Vcan, Megf11, Aqp4, Slc1a3, Gja1, Ebf1, Flt1) and the neural population separated out. The neural subset was then reclustered and neuron type identified using a separate list of markers (Syt1, Gad1, Gad2, Slc17a7, Ppp1r1b, Foxp2, Grm8, Drd1, Pdyn, Tac1, Ebf1, Drd2, Penk, Adora2a, Drd3, Elavl2, Chat, Kit, Pvalb, Lhx6, Sst, Vip, Calb1). Any clusters that had poor neural expression were removed and the resulting object reclustered at a resolution of 0.5 on 35 principle components. Common cell type subclusters were combined for analyses (e.g., D1-MSN).

Cell–cell communication networks were inferred by computing the probability of ligand–receptor interactions between cell types using CellChat (version 2.2.0) with default parameters. The resulting communication networks were aggregated to quantify the total number of inferred interactions and overall interaction strength for each condition. To enable direct comparison between conditions, CellChat objects were merged, and differences in network properties were assessed using built-in visualization and analysis functions, including circle plots, heatmaps, and network summary statistics. These metrics represent model-based inference of communication probability rather than direct statistical comparisons.

### Photometry Data Acquisition

A RZ10x with 405 (isosbestic) and 465 (GCaMP) nm LEDs were used to record photometry (Tucker Davis Technologies, Alachua, FL). Rats were connected via a 400 μm, NA 0.57 fiberoptic rotary joint (Doric Lenses, Montreal, QC) coupled to a 400 μm subject cable. An optical filter (Doric Lenses) was used to both restrict excitation light spectra and separate emitted fluorescence into isosbestic and calcium-dependent signals. Excitation light from the RZ10x was delivered to the filter via 2 m, 200 μm, 0.5 NA fiberoptic cables, and emitted signals were returned to the photodetector via 2 m, 600 μm, 0.5 NA cables. Light power was titrated per rat to optimize signal quality, defined by clear, dissociable calcium-dependent transients in the raw trace. Output powers ranged from 15–100 µW at the end of the cable (attenuated to ∼70% of value at tip of probe), with the 405 nm isosbestic channel set to 20–30% of the 465 nm excitation power.

### Photometry Data Processing

The raw photometry signal from 405 and 465 channels was binned into 50 ms means using TDT’s python-based script. Individual events (e.g., cue presentation, lever press) time stamps were then merged into these data in R where the remaining steps were performed. Quality control was then performed to remove disconnections and extreme signal changes that were not biologically plausible (e.g., cable fold due to rearing) based on visual inspection of data. A double exponential decay function was then fit to the target and isosbestic signals [P+(Y0-P)*(PctFast*exp(−Rate_fast*t) + (1-PctFast)*exp(−Rate_slow*t))], where P is the signal plateau, Y0 the initial intercept, PctFast the proportion of signal allocated to the first decay portion, Rate_fast/Rate_slow the rates for the early and late portion, and t is time]. Data were then normalized by subtracting the fitted signal from the raw signal. Then the isosbestic was subtracted from the calcium signal to remove motion and autofluorescent effects. Signals were then z-scored across the entire session and subsetted to the time around the target events. From these, the max peak, latency to peak, time to decay back to baseline (defined as +0.5 Z score), and area under the curve were calculated for analysis.

### Statistical Analyses

All statistical analyses were performed in R using the *lme4*, *emmeans*, and *brms* libraries. A model comparison approach was taken to determine if Sex X Injury interactions added explanatory power to statistical models. The Akaike Information Criterion was used to compare models. Mixed effects models were used for data with repeated measures. Individual subject intercepts were used as random effects. For univariate behavioral data (Pavlovian Conditioning, Conditioned Reinforcement, most RGT measures), electrophysiology data, histological data, and calcium photometry data linear regressions were used. For heavily-skewed and zero-inflated behavioral data on the RGT (impulsive responses, omissions), binomial regressions were used. For multivariate choice data on the RGT, a Bayesian linear mixed effects regression where choice option was also included in the random effect structure (ChoiceOption|Subject) was used to account for the interdependency [32]. Relations between behavioral tasks were evaluated with linear regressions.

Differential gene expression for single nucleus RNA sequencing data was performed using a pseudobulk method (*DESeq2* [64]) to control for false positives due to the large number of cells which would be treated as independent observations [65]. Differential gene expression was then analyzed at the aggregate level across all cells, and then for each individual cell cluster. A minimum threshold of *p* = 0.05 and absolute *z* score above 0.5 was considered significant. Significantly-affected genes were then loaded into Ingenuity Pathway Analysis (Qiagen USA, Germantown, MD) to evaluate affected pathways. The dominant signaling sources and targets of outgoing and incoming interaction strength between neuronal populations were quantitively inferred and analyzed using *CellChat* (v.1.6.1 [38]). The calculations were conducted using existing gene expression parameters and Seurat objects, with default settings. The CellChatDB mouse was used for analysis.

## Acknowledgements

Special thanks to Vonder Haar lab members who helped conduct behavioral testing, Brian Davis for assistance and feedback on nuclei extraction, fixation and barcoding, Andrew Fischer and Jeremy Day for feedback on sequencing processing, Myles Billard for assisting in development of calcium processing pipelines, and Miranda Koloski and Morteza Salimi for consultation on head cap surgeries in the TBI model. This work was funded by the NINDS (R01-NS110905 to CV), DOD (HT9425-23-1-1003 to CV, JPG, CCA), the International Center for Responsible Gaming (grant to CV) and OSU’s Center for Brain Injury Research and Discovery (summer undergraduate research fellowships to JEM & MAE). Work was supported by Ohio State University’s Genomics Core (NCI P30CA016058). Viruses were a gift from Karl Deisseroth (ChR2, Addgene viral prep #26976-AAV9) and James M. Wilson (GCaMP6f, Addgene viral prep #100834-AAV9).

## Author Contributions

Conceptualization – CV, EC, JEM; Formal analysis – CV, EC, SJ, PS, MP, JMP; Funding acquisition – CV, JPG, CCA; Investigation – EC, JEM, MAE, SJ, AGD, FBA, KMM, FZ; Visualization – EC, SJ, JEM, CV, PS; Writing – original draft – CV, EC, JEM; Writing – review & editing – CV, EC, JEC, AGD, FBA, KMM, PS, MP, CCA, JPG

## Notes

### Competing Interest Statement

The authors have declared no competing interest.

