## Supplement 1 for "Frontal Brain Injury Reduces Sensitivity to Reward-Predictive Cues and Remodels the Nucleus Accumbens"

### Supplemental Methods

Fiberoptic and Combined TBI/Fiberoptic Implant Surgery

The main methods present our final approach to this surgery. However, substantial piloting was required to establish a headcap that would last for months with a large open craniectomy. Modification was necessary to support repeated behavioral testing and chronic fiberoptic implantation. In initial surgeries, a fibrotic growth occurred under the headcap, resulting in progressive skull weakening – particularly surrounding burr holes, which ultimately resulted in implant instability and headcap loss. This issue was mitigated by fully sealing all exposed openings. A thin UV-cured cement layer was applied to cover the craniectomy and adjacent skull margins, and thin Metabond was used to flood and seal the screw/skull interfaces before building the headcap with acrylic-based cement. Long-term stability also depended critically on matching drill bit diameter precisely to screw size. Undersized burr holes increased insertion pressure and weakened surrounding bone, whereas oversized holes allowed fibrotic growth and implant loosening. Because procedures required extended surgical durations, maintaining skull hydration during preparation was essential to preserve bone integrity. However, hydration had to be carefully balanced with maintaining sufficiently dry surfaces during cement curing to avoid air gaps and bonding failure. Additional stability was achieved by positioning screws in the thickest available skull regions and as far as possible from the craniectomy margins.

Ultimately, sham procedures were “intact” with no craniectomy because sham animals had a higher risk of bleeding post-craniectomy from the mid-sagittal sinus (compared to TBI animals which experienced some swelling that halted bleeding). In the presence of a sealed headcap, such bleeding posed an elevated risk of intracranial pressure accumulation and required euthanasia. Therefore, sham surgeries were performed without craniectomy in the final protocol. An example surgical sheet can be requested by emailing the authors.

### Supplemental Results

Comparison of TBI data with Cocaine Transcriptome Data

In a prior publication, Brida and colleagues (2025, *Sci Advances*) identified reelin as a driver of responsiveness to cocaine in NAc neurons. Given prior data showing vulnerability to cocaine self-administration in TBI (Vonder Haar et al., 2019, *Eur J Neurosci*), we decided to directly evaluate the reelin gene *Reln* in the current data.

In the current data, reelin was high in the D1 and mixed MSN populations relative to other MSNs. It was not significantly differential in pseudobulk transcriptional analyses. However, when plotted and analyzed at the cell transcript level, TBI significantly reduced *Reln* expression in the D1-, D2-, and mixed-MSN cell types ($p$’s < 0.001; Fig S12). These data suggest that other cell types are more likely to drive vulnerability to substance abuse in TBI.

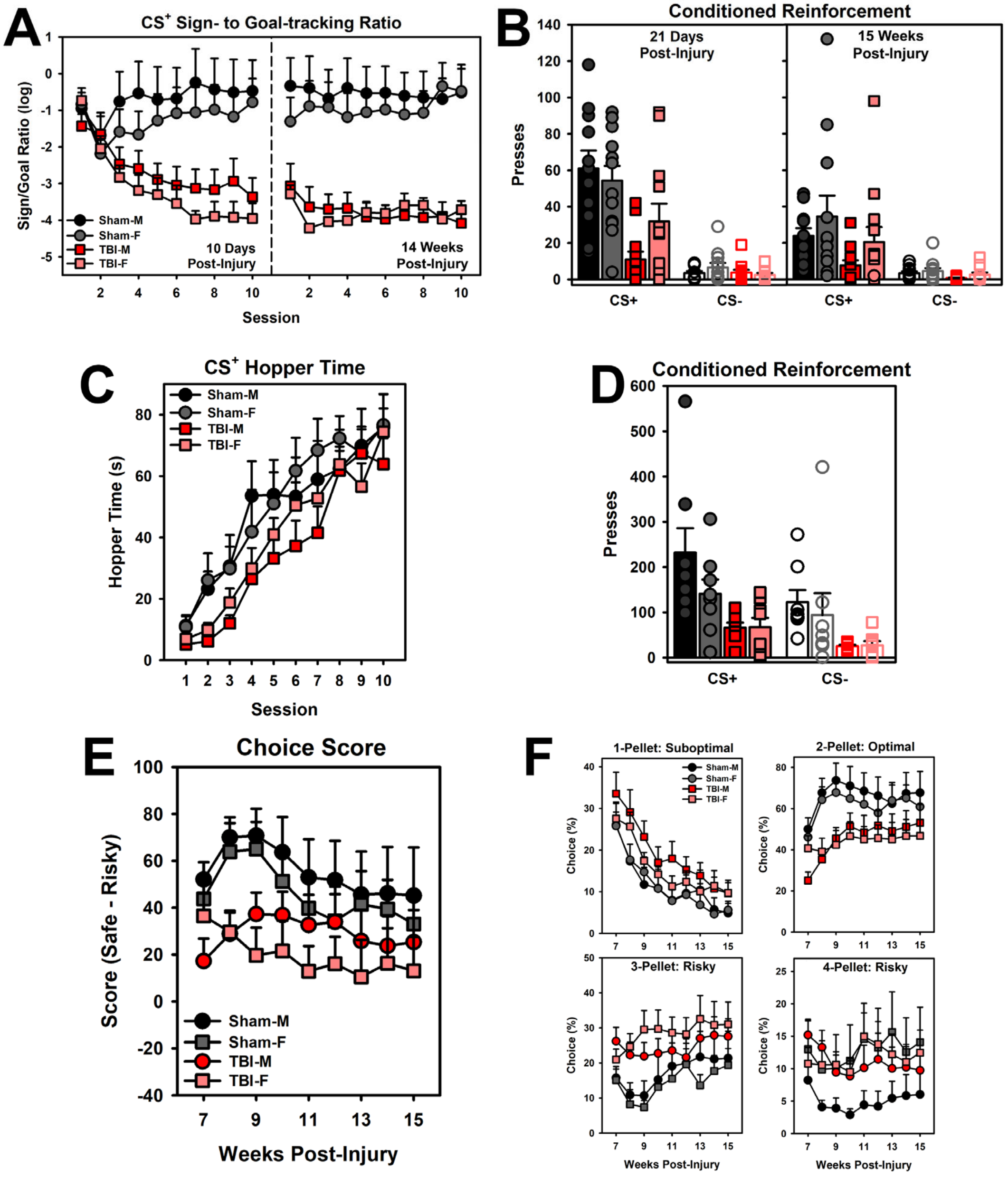


**Figure S1. Breakdown of male/female data corresponding to manuscript Figure 1.** Interacting sex with injury did not improve the model for any variables. A) For visual conditioned reinforcement, there was no main effect of sex on acute or chronic sign-to-goal tracking ratio ($p$'s > 0.282). B) For visual conditioned reinforcement, models with sex included did not improve the overall fit, suggesting no effect of sex. C) For auditory conditioned reinforcement, there was no main effect of sex goal tracking behavior ($p$ = 0.460). D) For auditory conditioned reinforcement, models with sex included did not improve the overall fit, suggesting no effect of sex. E) There was no main effect of sex on RGT Choice Score ($p$ = 0.353). F) The breakdown of the choices showed no main effect of sex (CI: -0.02 to +0.02).

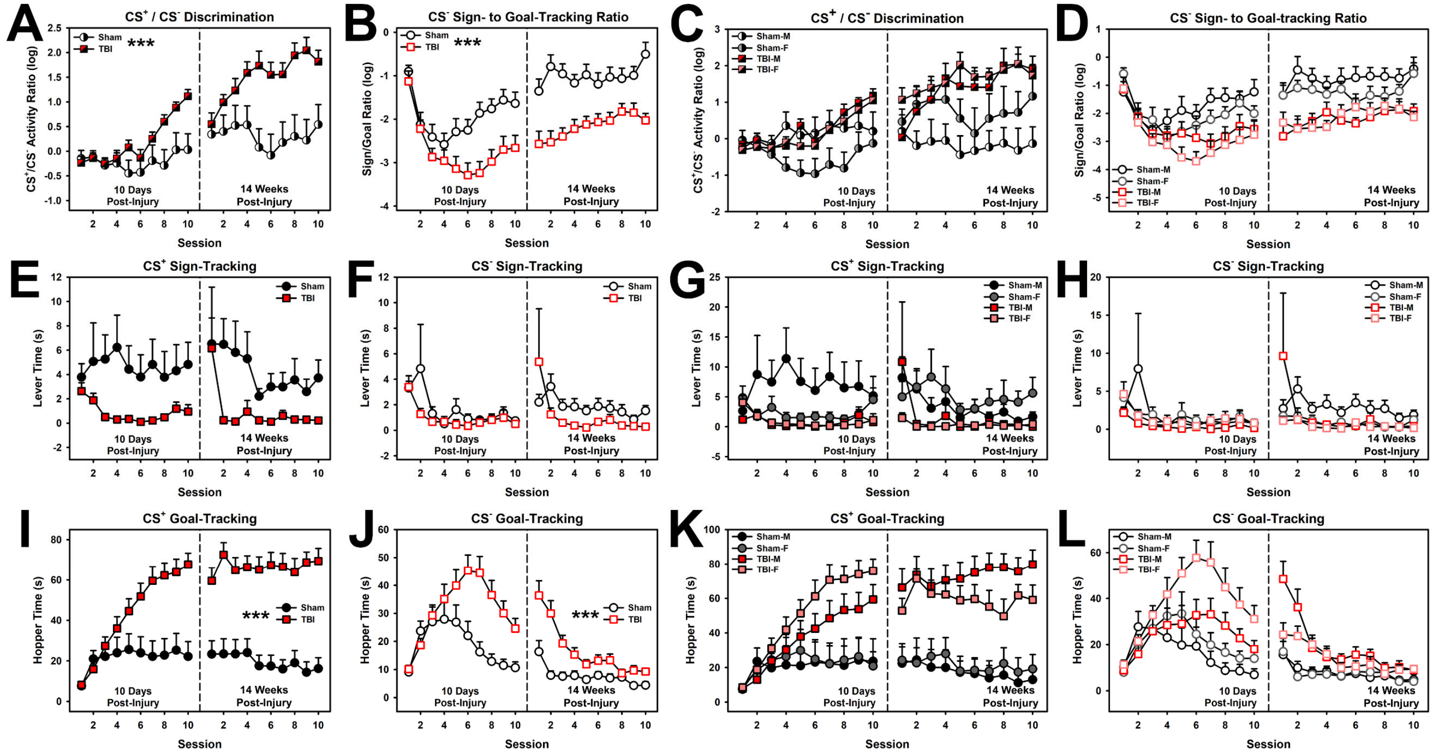


**Figure S2. Other variables captured during visual Pavlovian conditioning and their breakdown by sex.** Adding sex by injury interactions to models did not improve their fit, suggesting no interaction with injury. A) There was an interaction between trial type, session, and injury ($p < 0.001$) such that TBI rats showed more activity during CS+ presentations relative to CS- presentation (primarily driven by a drop-off in high goal-tracking behaviors during CS- [see panel J]). B) Sign- vs. goal-tracking during CS- cue presentation. TBI rats showed less sign-tracking to the CS- than sham ($p < 0.001$). C) Sex breakdown of data in panel A. There was no main effect of sex ($p = 0.232$). D) Sex breakdown of data in panel B. There was no main effect of sex ($p = 0.213$). E) Sign-tracking during CS+ cue presentation. There was no effect of injury or injury by time interaction ($p$'s > 0.099). F) Sign-tracking during CS- cue presentation. There was no effect of injury or injury by time interaction ($p$'s > 0.149). G) Sex breakdown of data in panel E. There was no main effect of sex ($p = 0.425$). H) Sex breakdown of data in panel F. There was no main effect of sex ($p = 0.233$). I) Goal-tracking during CS+ cue presentation. Injury substantially increased goal-tracking behavior ($p < 0.001$). There was no effect of injury or injury by time interaction ($p$'s > 0.099). J) Goal-tracking during CS- cue presentation. Injury substantially increased goal-tracking behavior, though it eventually trended back down for the CS- ($p < 0.001$). K) Sex breakdown of data in panel I. There was no main effect of sex ($p = 0.333$). L) Sex breakdown of data in panel L. There was a main effect of sex, with males showing less goal tracking ($p = 0.034$). *** = $p < 0.001$.

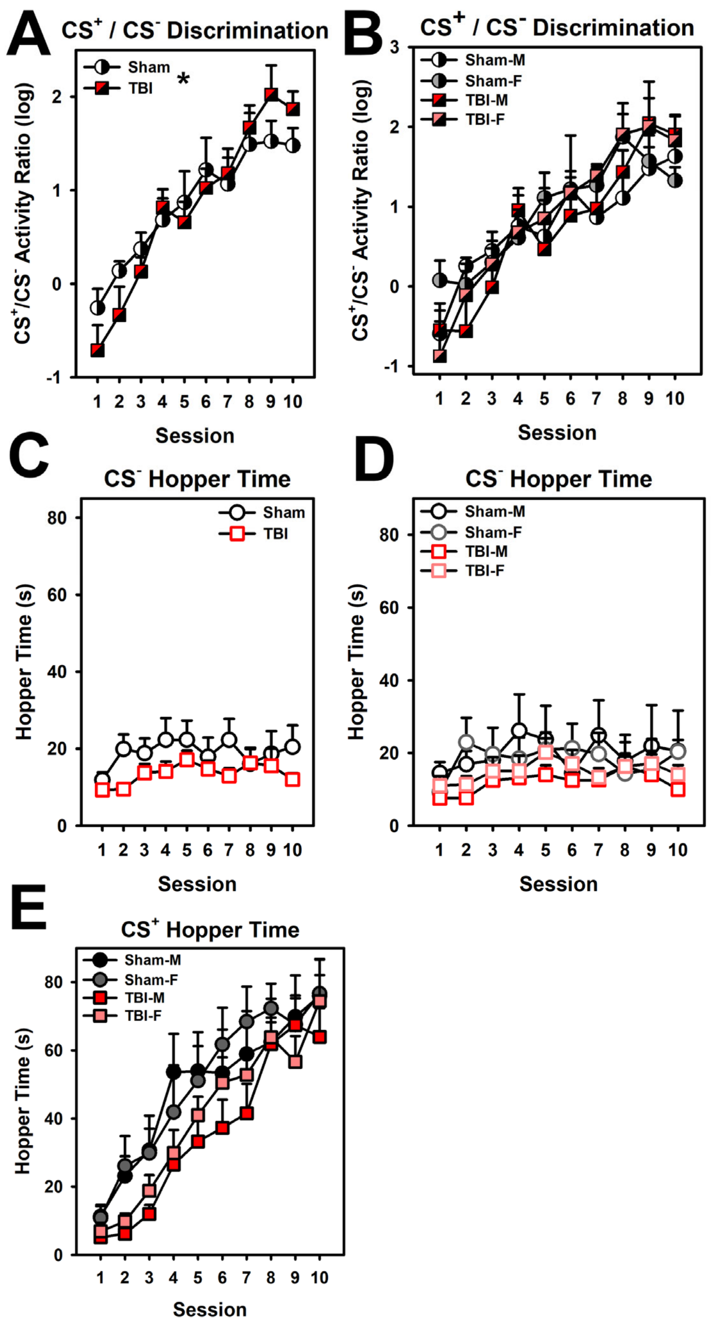


**Figure S3. Other variables captured during auditory Pavlovian conditioning and their breakdown by sex.** Adding sex by injury interactions to models did not improve their fit, suggesting no interaction with injury. A) There was an interaction between injury and session ($p$ = 0.012) such that TBI rats started slightly lower and increased faster relative to shams in discriminating CS+ from CS- presentations. B) Sex breakdown of data in panel A. There was no main effect of sex ($p$ = 0.652). C) Goal-tracking during CS- cue presentation. There was no significant difference ($p$ = 0.774). D) Sex breakdown of data in panel C. There was no main effect of sex ($p$ = 0.867). E) Sex breakdown of goal-tracking during CS- cue presentation (corresponding to main Figure 1E). There was no significant main effect of sex ($p$ = 0.460). * = $p$ < 0.005.

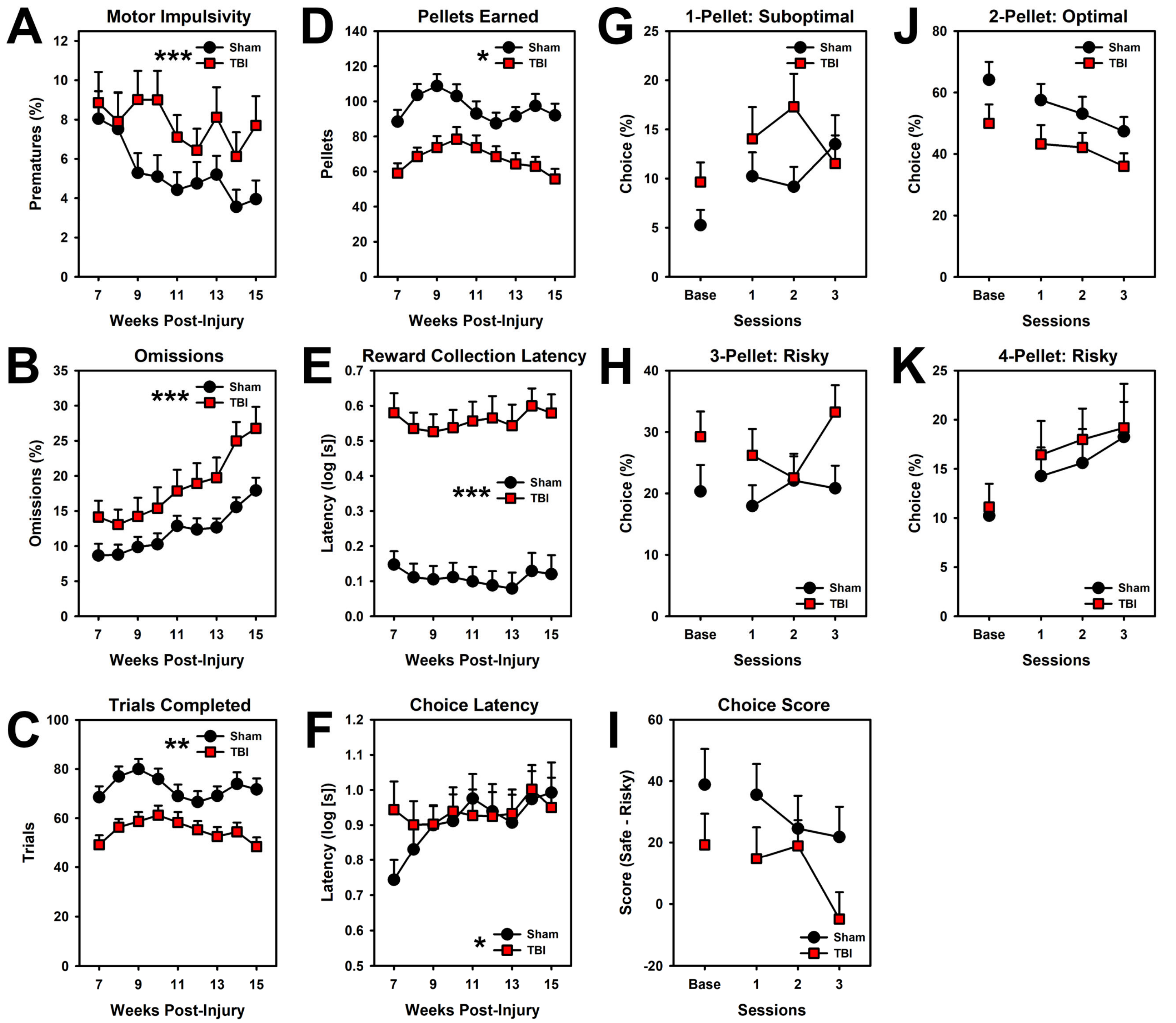


**Figure S4. Other variables captured during cRGT.** Adding sex by injury interactions to models did not improve their fit, suggesting no interaction with injury. A) Impulsivity as measured by premature responses. TBI increased impulsivity over time ($p < 0.001$). B) TBI increased omissions over time ($p < 0.001$). C) TBI reduced trials completed ($p = 0.009$). D) TBI reduced pellets earned over time ($p = 0.012$). E) TBI increased reward collection latency ($p < 0.001$). F) TBI started higher and did not increase their duration to make choices ($p = 0.023$). G-K) Choice of the various options during extinction. Comparison of Bayesian models indicated that TBI did not improve model fit, suggesting no effect of injury on rate of extinction. I) Score variable summary of choice showing no effect of injury in change from baseline choice ($p = 1.0$). *** = $p < 0.001$, ** = $p < 0.01$, * = $p < 0.005$.

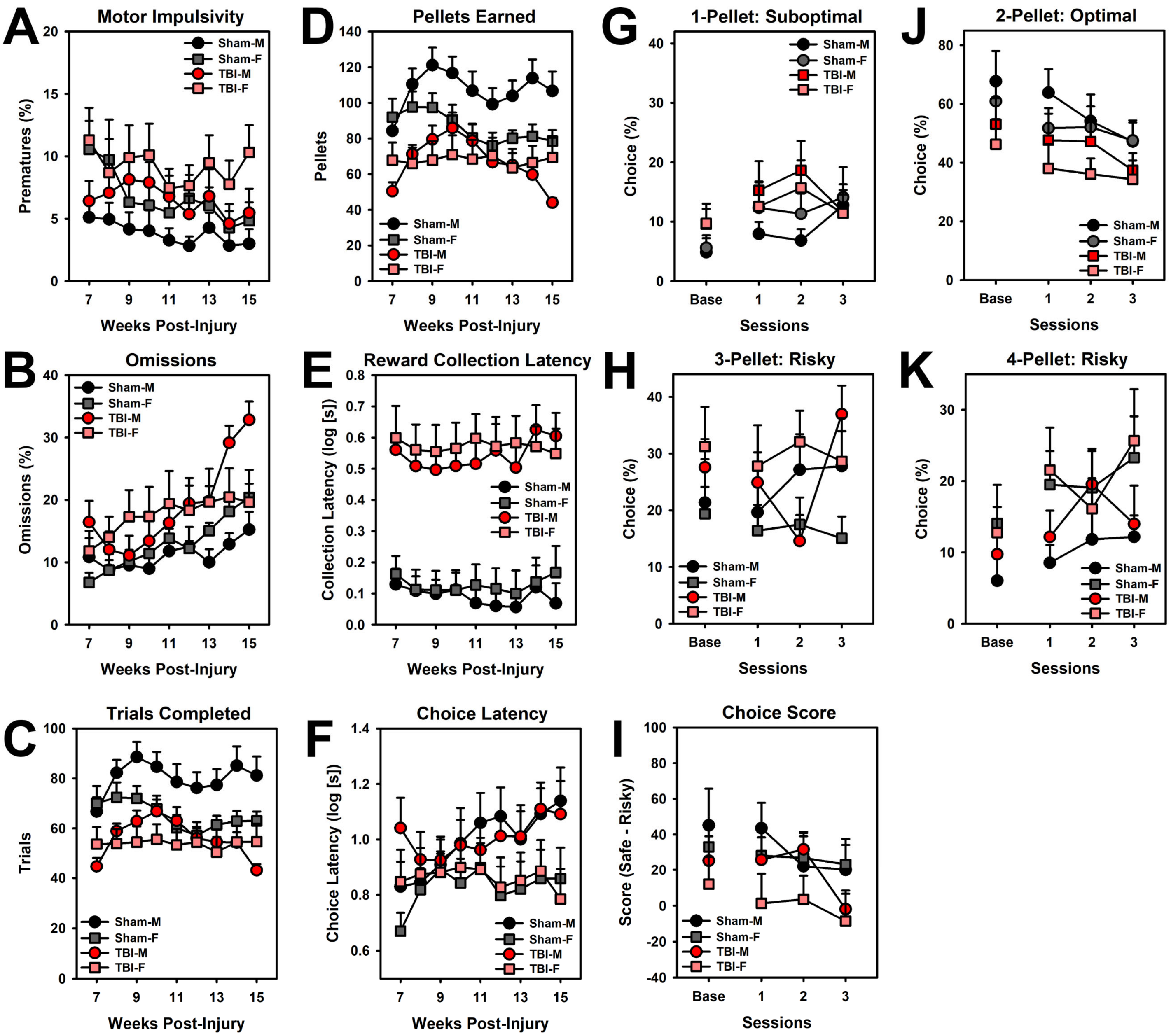


**Figure S5. Breakdown of male/female data corresponding to Figure S4.** Interacting sex with injury did not improve the model for any variables. A) There was no main effect of sex ($p$ = 0.078). B) There was no main effect of sex ($p$ = 0.705). C) There was no main effect of sex ($p$ = 0.061). D) There was no main effect of sex ($p$ = 0.142). E) There was no main effect of sex ($p$ = 0.517). F) There was no main effect of sex ($p$ = 0.100). G-K) Comparison of Bayesian models indicated that sex did not significantly improve the model fit, suggesting no effect of sex. I) There was no main effect of sex ($p$ = 0.353).

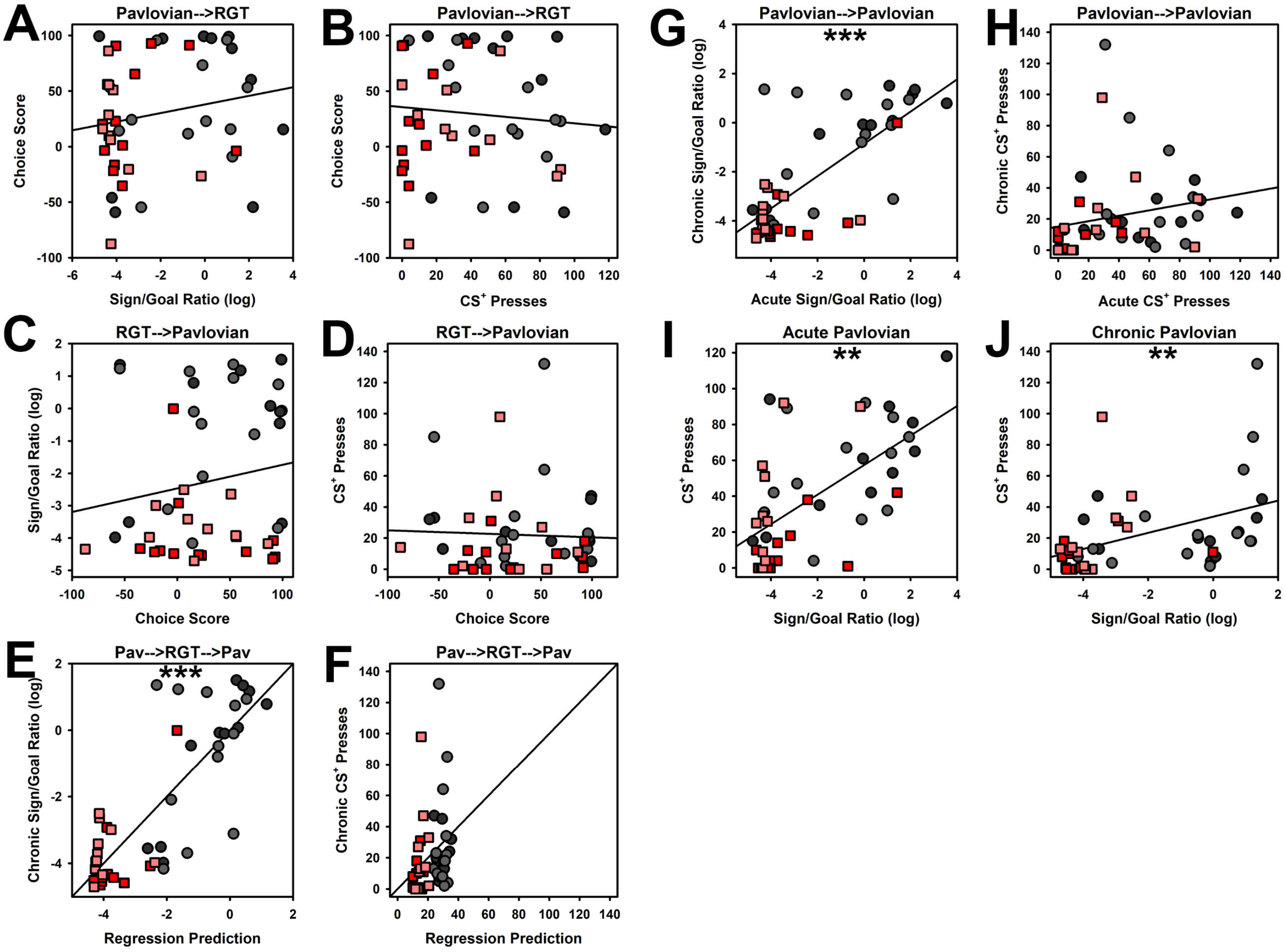


**Figure S6. Correlations between behavioral measurements including Pavlovian conditioned approach (Sign/Goal-tracking), conditioned reinforcement (presses), RGT choice (Score).** Models interacting injury were never significantly better than those with only main effects of injury, suggesting that injury did not modify any task relations. A) Initial sign/goal-tracking ratio did not predict RGT choice ($p = 0.212$). B) Initial conditioned reinforcement did not predict RGT choice ($p = 0.138$). C) RGT choice did not predict subsequent sign/goal-tracking ratios ($p = 0.677$). D) RGT choice did not predict subsequent conditioned reinforcement ($p = 0.527$). E) Initial sign/goal-tracking ratio predicted subsequent sign/goal-tracking ratios ($p < 0.001$), but was not modified by RGT choice ($p = 0.784$). F) Initial conditioned reinforcement did not predict subsequent conditioned reinforcement ($p = 0.696$) and was not modified by RGT choice ($p = 0.970$). G) Initial sign/goal-tracking ratios predicted subsequent sign/goal-tracking ratios ($p < 0.001$). H) Initial conditioned reinforcement did not predict subsequent conditioned reinforcement ($p = 0.151$). I) Sign/goal-tracking ratios predicted conditioned reinforcement presses during the acute testing phase ($p = 0.002$). J) Sign/goal-tracking ratios predicted conditioned reinforcement presses during the chronic testing phase ($p = 0.003$). Symbols indicate individual rats; lighter colors are females; *** = $p < 0.001$, ** = $p < 0.01$.

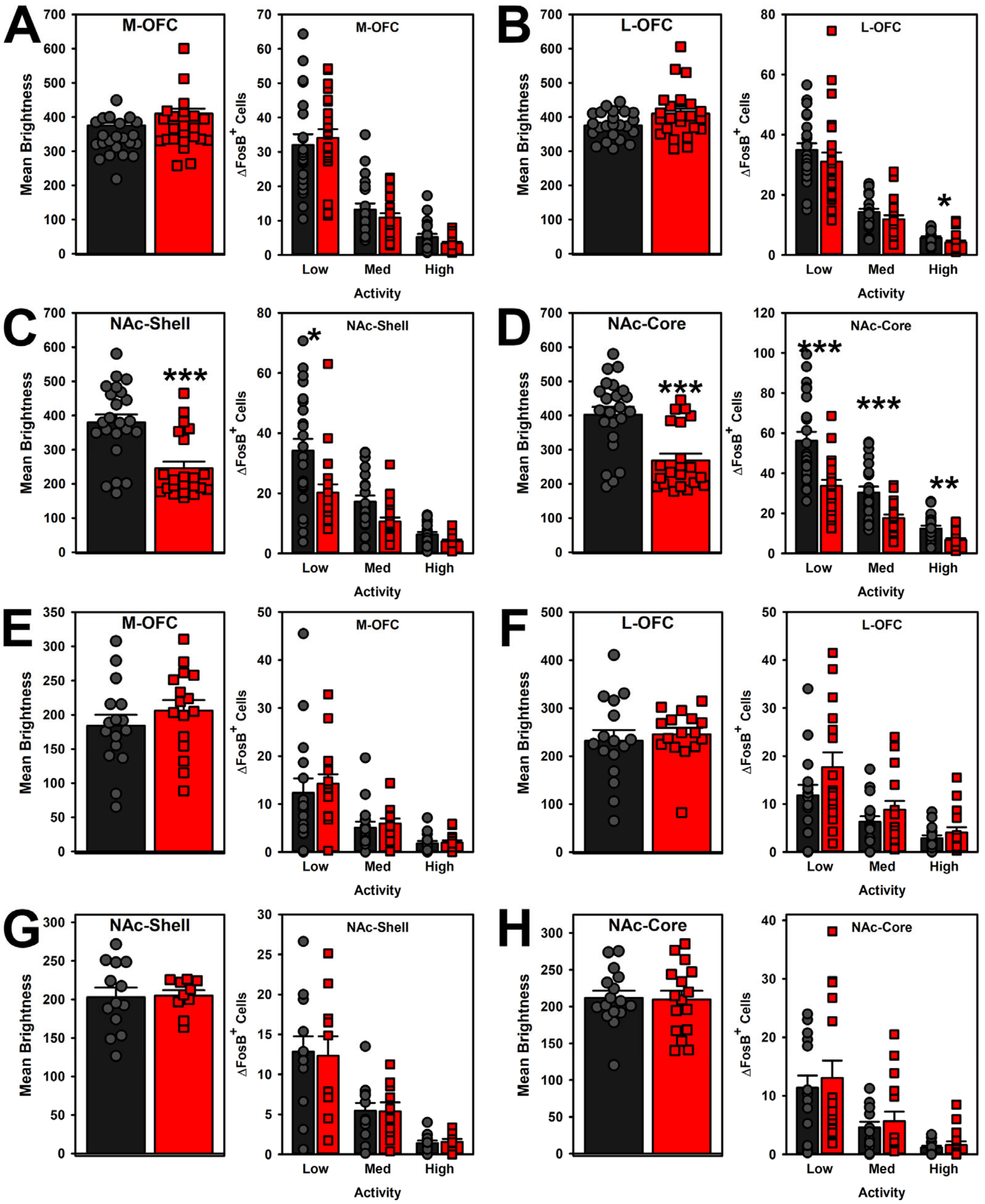


**Figure S7. Additional data on mean brightness and counts thresholded by brightness/activity supporting main figure A-C.** Note that "low" activity bars are what are reported in the main manuscript and are reproduced here for ease of comparison. A) M-OFC brightness was not significantly affected by injury ($p$ = 0.053). M-OFC cell counts were not changed by the injury at low, medium, or high thresholds ($p$'s > 0.230). B) L-OFC brightness was not significantly affected by injury ($p$ = 0.722). Injury did not affect cell counts at the low or medium activity thresholds ($p$ = 0.135; $p$ = 0.064), however, when the most active cells were thresholded, TBI reduced counts ($p$ = 0.012). C) TBI decreased brightness for the NAc Shell ($p$ < 0.001). Injury reduced cell counts at the lowest activity threshold ($p$ = 0.016), however this effect faded at medium ($p$ = 0.051) and high activity thresholds ($p$ = 0.109). D) TBI decreased brightness for the NAc Core ($p$ < 0.001). TBI decreased counts at low, medium, and high activity thresholds ($p$ < 0.001; $p$ < 0.001; $p$ = 0.003). E-H) TBI had no effect on any regions when measured at 22 days post-injury after only 10 days of auditory Pavlovian conditioning (p's > 0.118). *** = $p$ < 0.001, ** = $p$ < 0.01, * = $p$ < 0.005.

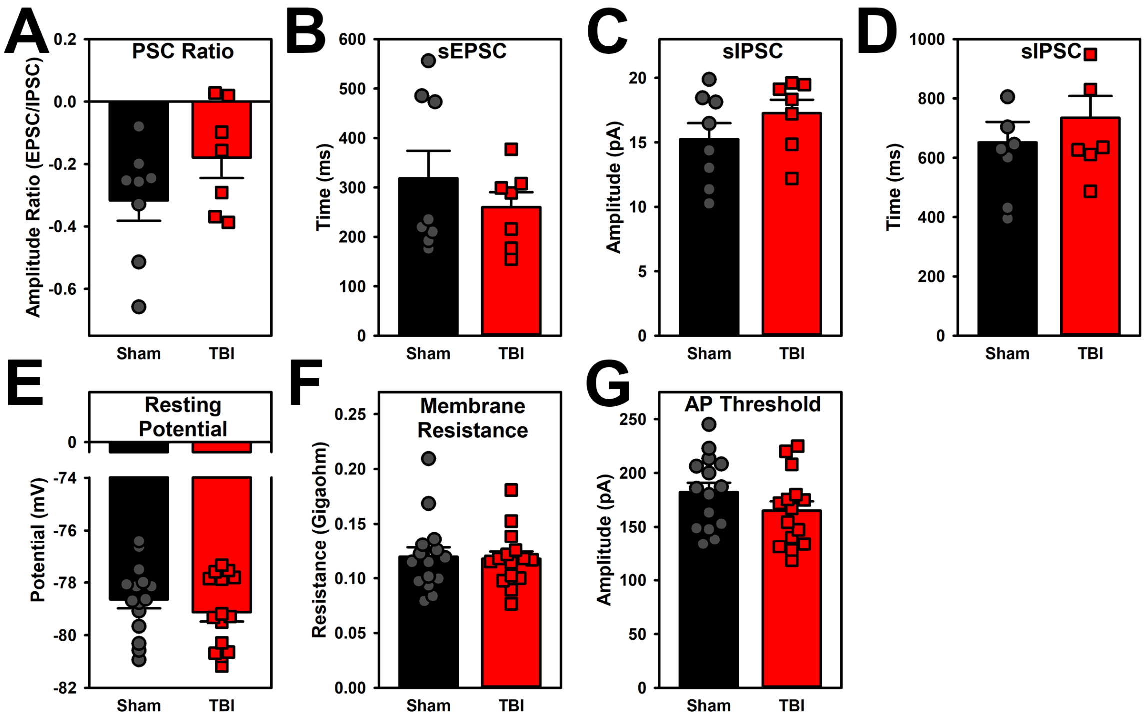


**Figure S8. Cell properties during patch-clamp recordings.** A) The postsynaptic current ratio (EPSC/IPSC amplitude) was not affected by injury ($p$ = 0.231). B) The mini excitatory postsynaptic current interval was not affected by injury ($p$ = 0.375). C) The miniature inhibitory postsynaptic current amplitude was not affected by injury ($p$ = 0.162). D) The miniature inhibitory postsynaptic current interval was not affected by injury ($p$ = 0.438). E) The resting membrane potential was not affected by injury ($p$ = 0.268). F) The neuron's resistance was not affected by injury ($p$ = 0.826). G) Despite an increasing rate of spikes to stimulation (main Figure 2F), the minimum threshold for the first action potential was not significantly different ($p$ = 0.142).

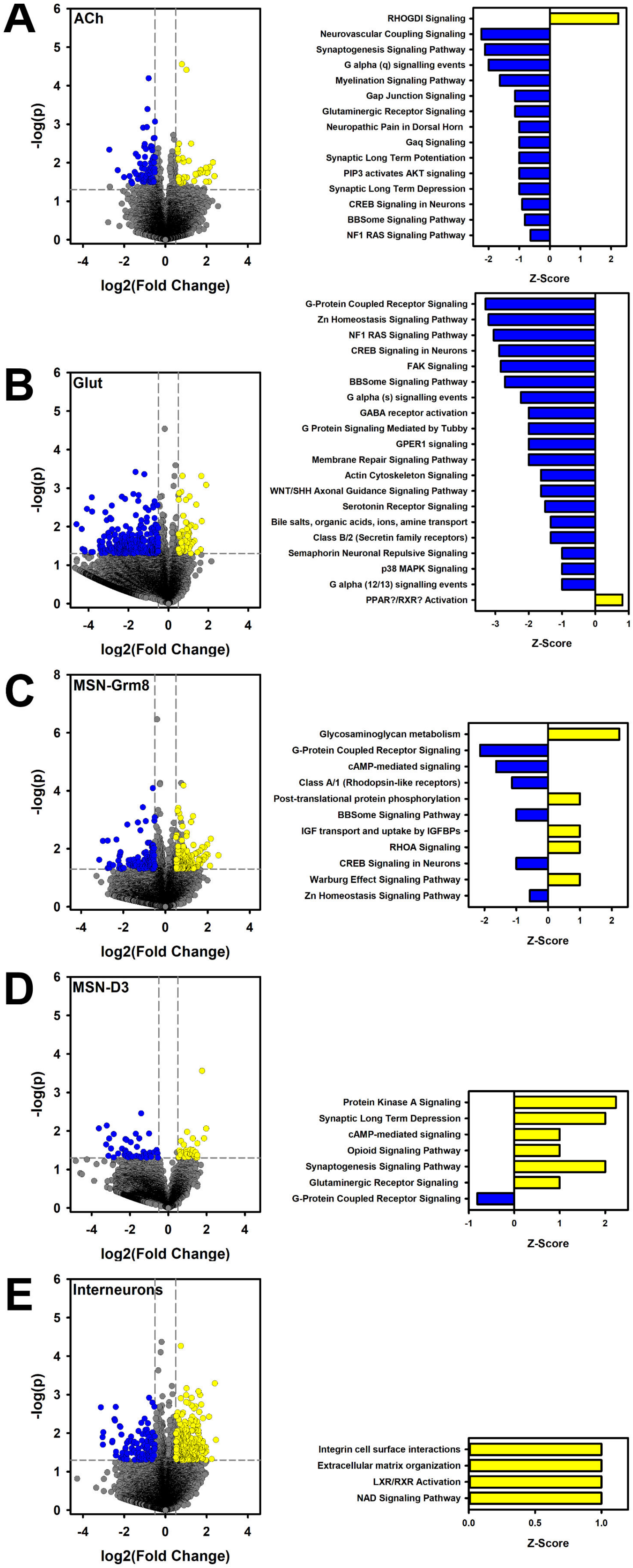
A
ACh
-log(p)
log2(Fold Change)
RHOGDI Signaling
Neurovascular Coupling Signaling
Synaptogenesis Signaling Pathway
G alpha (q) signalling events
Myelination Signaling Pathway
Gap Junction Signaling
Glutaminergic Receptor Signaling
Neuropathic Pain in Dorsal Horn
Gaq Signaling
Synaptic Long Term Potentiation
PIP3 activates AKT signaling
Synaptic Long Term Depression
CREB Signaling in Neurons
BBSome Signaling Pathway
NF1 RAS Signaling Pathway
Z-Score
B
Glut
-log(p)
log2(Fold Change)
G-Protein Coupled Receptor Signaling
Zn Homeostasis Signaling Pathway
NF1 RAS Signaling Pathway
CREB Signaling in Neurons
FAK Signaling
BBSome Signaling Pathway
G alpha (s) signalling events
GABA receptor activation
G Protein Signaling Mediated by Tubby
GPER1 signaling
Membrane Repair Signaling Pathway
Actin Cytoskeleton Signaling
WNT/SHH Axonal Guidance Signaling Pathway
Serotonin Receptor Signaling
Bile salts, organic acids, ions, amine transport
Class B/2 (Secretin family receptors)
Semaphorin Neuronal Repulsive Signaling
p38 MAPK Signaling
G alpha (12/13) signalling events
PPAR?/RXR? Activation
Z-Score
C
MSN-Grm8
-log(p)
log2(Fold Change)
Glycosaminoglycan metabolism
G-Protein Coupled Receptor Signaling
cAMP-mediated signaling
Class A/1 (Rhodopsin-like receptors)
Post-translational protein phosphorylation
BBSome Signaling Pathway
IGF transport and uptake by IGFBPs
RHOA Signaling
CREB Signaling in Neurons
Warburg Effect Signaling Pathway
Zn Homeostasis Signaling Pathway
Z-Score
D
MSN-D3
-log(p)
log2(Fold Change)
Protein Kinase A Signaling
Synaptic Long Term Depression
cAMP-mediated signaling
Opioid Signaling Pathway
Synaptogenesis Signaling Pathway
Glutaminergic Receptor Signaling
G-Protein Coupled Receptor Signaling
Z-Score
E
Interneurons
-log(p)
log2(Fold Change)
Integrin cell surface interactions
Extracellular matrix organization
LXR/RXR Activation
NAD Signaling Pathway
Z-Score

**Figure S9. Volcano plots breaking down data presented in table from main Fig 4A and corresponding pathway loadings.** A-B) Cholinergic and glutamatergic cells largely showed downregulation of many plasticity-related and cellular signaling-related pathways as a potential counterbalance to the large-scale change in the inhibitory cells. C) Grm8-MSNs showed the most mixed response to injury of the MSNs. Several intracellular signaling pathways were downregulated. D) The small D3-MSN class showed a similar but attenuated effect relative to the D1- and D2-MSNs. E) The interneurons had substantial gene changes but these mapped to very few pathways. A breakdown of cell types showed similar gene changes across the various subclasses (see Supplement 2).

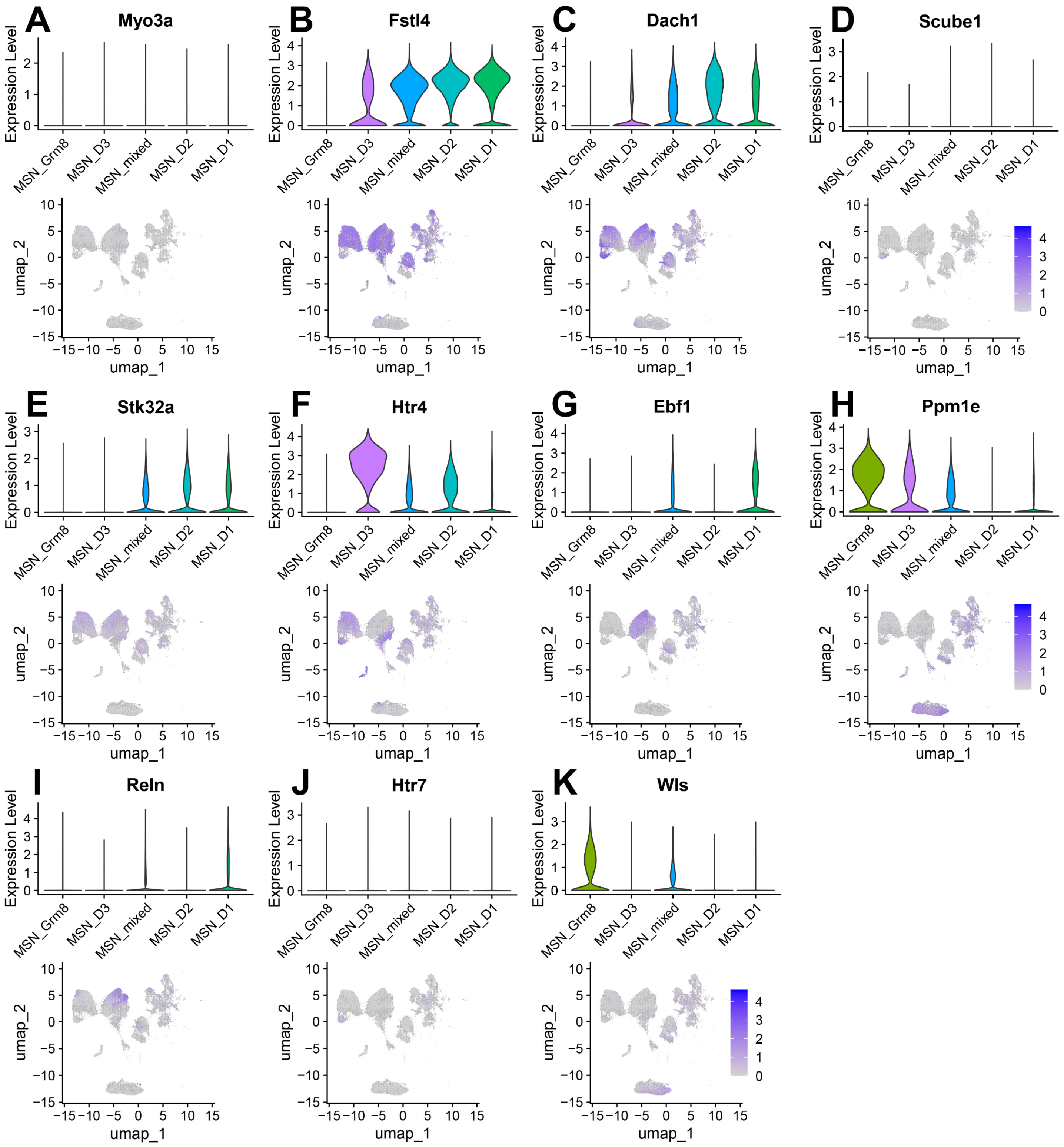


**Figure S10. Gene profile of potential marker genes of D1 and D2 MSNs from Reiner et al., 2024, *Sci Reports* and Fitzgerald et al., 2023, *eBioMedicine*.** The MSN-mixed phenotype showed elements of D1-, D2-, and even Grm8-MSN cell types. A-K) Violin plots of individual genes and their localization to clusters.

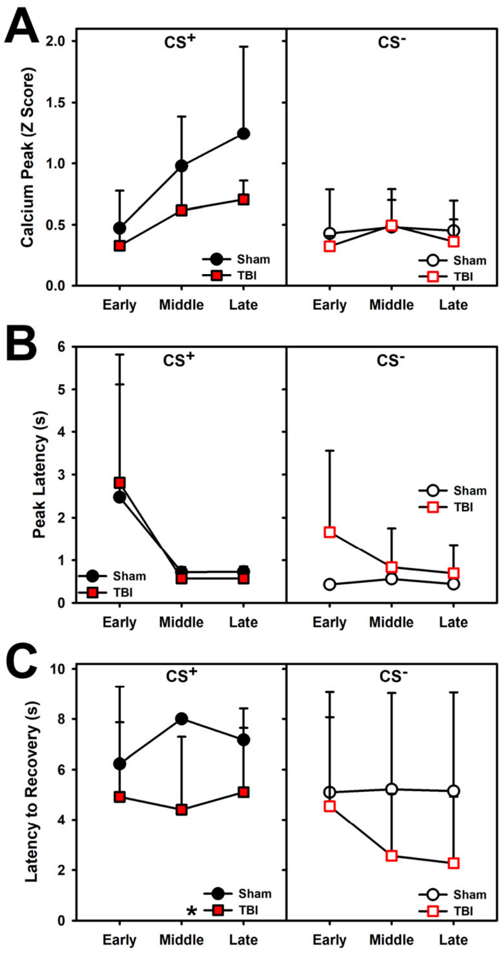


**Figure S11. Additional variables quantifying data presented in main Figure 5C-D.** A) Peak of calcium traces were not significantly affected by injury for CS+ or CS- (*p*'s > 0.084). B) Latencies for calcium peak to occur were not significantly affected by injury for CS+ or CS- (*p*'s > 0.403). C) The latency for the peak to decay was significantly reduced by injury for the CS+ (*p* = 0.021), but not significant for the CS- (*p*'s > 0.230). * = *p* < 0.005.

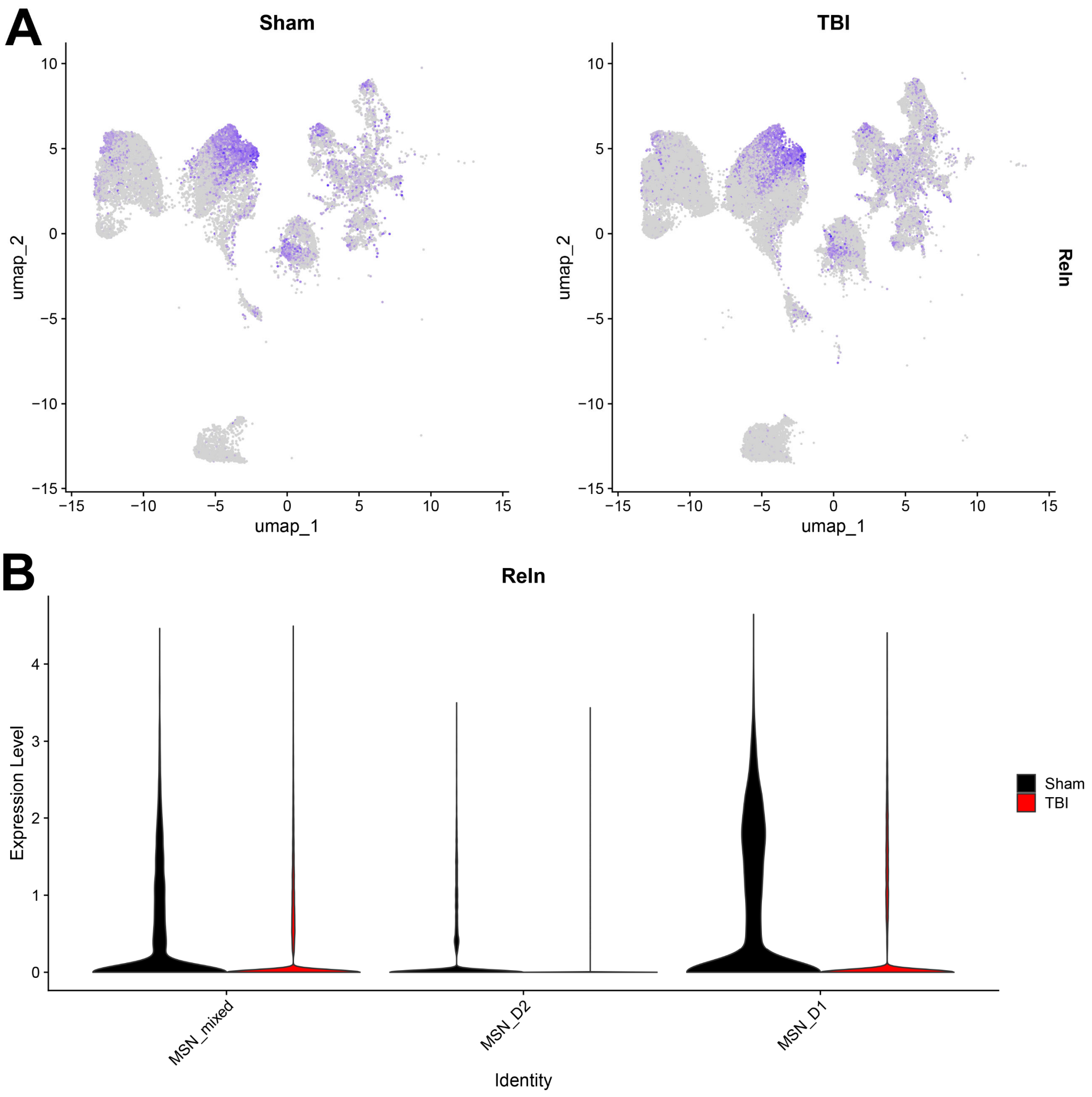


**Figure S12. Expression of the reelin gene *Reln* after TBI in D1-, D2-, and mixed-MSN cell types.** A) Expression plot overlaid on UMAP. B) Violin plots demonstrating that TBI significantly reduced expression in D1-MSNs (log2FC = 1.00, $p < 0.001$), D2-MSNs (log2FC = 0.54, $p < 0.001$), and mixed-MSNs (log2FC = 1.82, $p < 0.001$).
